# Hepatitis B virus neutralization with DNA origami nanoshells

**DOI:** 10.1101/2023.12.20.572526

**Authors:** Elena M. Willner, Fenna Kolbe, Frank Momburg, Ulrike Protzer, Hendrik Dietz

## Abstract

We demonstrate the use of DNA origami to create virus-trapping nanoshells that efficiently neutralize hepatitis B virus (*HBV*) in cell culture. By modifying the shells with a synthetic monoclonal antibody that binds to the *HBV* envelope, the effective neutralization potency per antibody is increased by approximately 100 times compared to using free antibodies. The improvements in neutralizing the virus are attributed to two factors: first, the shells act as a physical barrier that block the virus from interacting with host cells; second, the multivalent binding of the antibodies inside the shells lead to stronger attachment to the trapped virus, a phenomenon known as avidity. Pre-incubation of shells with *HBV* and simultaneous addition of both components separately to cells lead to comparable levels of neutralization, indicating rapid trapping of the virions by the shells. Our study highlights the potential of the DNA shell system to rationally create novel antivirals using components that, when used individually, show little to no antiviral effectiveness.

---

Programmable self-assembly with DNA origami allows the creating of user-defined 3-dimensional nanoscale structures. These structures can be used for a wide range of applications, including drug delivery, biosensing, molecular motors, and as templates for synthesizing inorganic materials.^1-6^ The programmability of DNA origami enables precise control over size, shape, and composition, making it a versatile platform for constructing nanoscale structures with tailored functionalities. In DNA origami, sets of DNA single-strands (“staples”) are designed to base pair with a long single-stranded DNA molecule (“scaffold”) to fold it into a predefined shape.^7-9^ Multiple discrete DNA origami building blocks, in turn, can then oligomerize into well-defined higher order 3D objects^10^ with dimensions that can exceed those of viruses^11^, and it has been explored using such shells and other DNA nanoarchitectures to inhibit the entry of viruses to cells.^12-16^ In principle, any virus binder could be utilized to coat the inside of the DNA origami nanoshell. This platform is particularly useful as a virus entry inhibitor when employing moieties that bind to a target virus but lack substantial virus-neutralizing properties. In addition, the efficacy of virus binders with pre-existing neutralization capacity is expected to be enhanced when used in the context of the shells. This is because we expect the shell material to contribute to blocking viruses from interactions with cells by presenting a physical barrier to infection. Furthermore, the density of the virus binders inside the DNA shells may be controlled by the user so that avidity effects stemming from the simultaneous binding of multiple virus binders to the same virus can be elicited, which is also expected to enhance neutralization potency. In the present work, we test the neutralization capacity of DNA origami nanoshells, exemplarily in cell cultures using infectious and replicative hepatitis B viruses as a model system.

The hepatitis B virus (*HBV*) is an enveloped virus of the *hepadnaviridae* family. *HBV* predominantly infects hepatocytes and can cause significant liver damage resulting in liver cirrhosis, and is the single most frequent cause of liver cancer. This cancer is known as hepatocellular carcinoma, which is hard to treat and one of the most deadly cancers.^17, 18^ *HBV* is a spherical enveloped virus with an approximately 42 nm diameter (Figure 1A) containing an icosahedral capsid that encapsulates the viral DNA genome, which occurs in a relaxed circular form (*rcDNA*) and has a length of about 3200 bp.^19^ The capsid is a homo-polymer built from 180 or 240 subunits of the *HBV* core protein (*HBcAg*, Figure 1A, red). The *HBV rcDNA* genome is formed in the viral capsid by a DNA polymerase with a reverse-transcriptase activity using a 3.5 kb pre-genomic RNA as a template. The envelope consists of a lipid bilayer membrane densely packed with the virus surface proteins, the large (*L*), medium *(M*), and small (*S*) surface proteins, with the latter being produced in excess.

After infection of a hepatocyte, the *rcDNA* is transported to the nucleus and converted to the covalently closed-circular DNA (*cccDNA*), which is the nuclear persistence form of *HBV. HBV* replication in the cell produces an additional antigen known as the hepatitis B e antigen (*HBeAg*, Figure 1A, yellow). *HBeAg* is secreted by the host cell and can be used as a marker of active viral replication.^20^ In addition to mature *HBV* virions, *HBV*-positive cells typically also secrete noninfectious spherical and tubular subviral particles mainly consisting of the outer-envelope proteins *S*. These subviral particles are smaller than the actual *HBV* virions, with diameters of approximately 20-22 nm.^21^ Hepatitis B viruses fit into a previously described DNA nanoshell prototype^12^ (Fig. 1B). Na^+^-taurocholate co-transporting polypeptide (*NTCP*) expressing *HepG2* cells present a reliable in vitro system to quantify *HBV* infection^22, 23^ and infection neutralization^24^. Markers such as *HBeAg* and *cccDNA* can be used to analyze the level of *HBV* infection. We used a synthetic antibody (*MoMAb*) that consists of 2 copies of a single chain antibody fragment fused to a mutated fragment crystallizable (*Fc)* domain of an immunoglobulin G1 *(IgG1)* with reduced *Fc*-receptor binding affinity^25^ to coat DNA nanoshells and test their neutralization capacity relative to that of the free antibodies.

**Figure 1.**
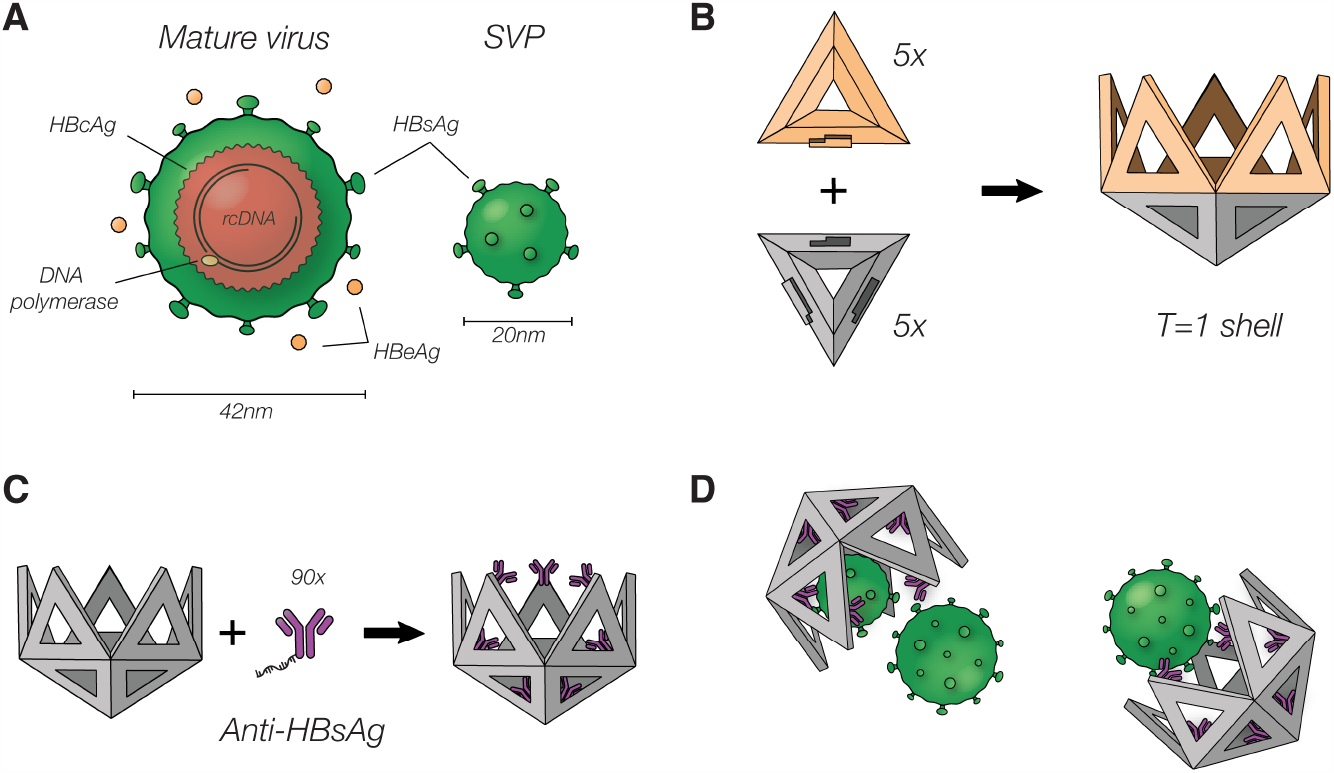
Hepatitis B overview and capture using DNA nanoshells. **(A)** Illustration of the Hepatitis B virus. Left the mature virus, right the subviral particle (SVP) consisting of solely *HBsAg* **(B)** Visualization of the T=1 nanoshell (right) and its 2 different triangular subunits (left). 5 gray triangles build the pentameric base and 5 yellow triangles bind to this base to create a deeper cavity. **(C)** Functionalization of the T=1 nanoshell: *HBsAg* antibodies are functionalized with a *ssDNA* strand and mounted to the 90 binding sites inside the shell. **(D)** Functionalized nanoshells can specifically recognize and bind *HBV* and its subviral particles.

## Results

To trap hepatitis B virus particles, we created icosahedral DNA origami nanoshells with an 80 nm wide cavity (Figure 1B), using 2 distinct triangular building blocks as previously described.^12^ To stabilize the shells for cell culture, we covalently crosslinked the constituent triangular subunits using UV point welding.^26^ We included 9 single-stranded DNA (*ssDNA*) handles that protrude from each of the triangular subunits, creating 90 attachment sites in the shell’s internal cavity. We selected a synthetic immunoglobulin G (*IgG*) antibody against *HBsAg*, referred to as *MoMAb*, as a binder,^25^ and labeled it with thiolated DNA single-strands that were complementary to the handles displayed on the interior surface of the shells. To tag the antibodies with DNA, we used a sulfosuccinimidyll 4-(N-maleimidomethyl) cyclohexan-1-carboxylat (*sulfo-SMCC*) crosslinker to target surface-exposed lysine residues on the *IgG* (Supplementary Figure 2, 3). We then added the DNA-tagged *MoMAb* in a 90:1 excess to a solution containing the DNA shells to populate all binding sites inside the shell cavity (Figure 1C). With this coating, we could successfully trap *HBV* in DNA nanoshells (Figure 1D).

**Figure 2.**
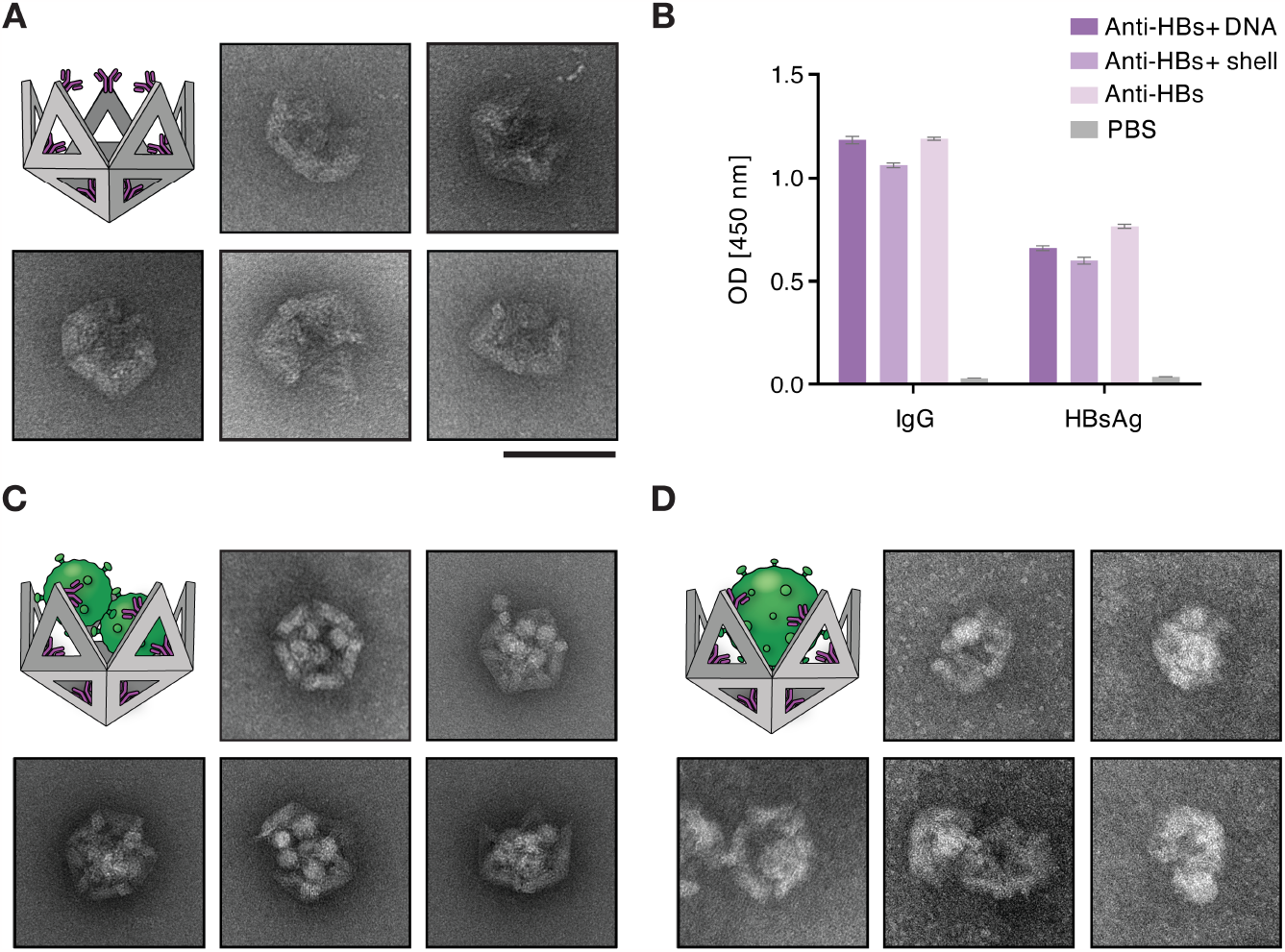
Shell functionalization and resulting *HBV* subviral particle and virion capture. **(A)** Negative stain *TEM* images of the functionalized DNA shell using the *anti-HBs* antibody *MoMAb*. **(B)** Binding control of the functionalized *anti-HBs* antibody. Nanoshells conjugated with *MoMAb anti-HBs* antibodies (medium violet) and with *anti-HBs* conjugated to the DNA handle without formation of nanoshells (dark violet) were analyzed by *ELISA* with either anti-*IgG* or *HBsAg* immobilized to the plates. *MoMAb* alone (light violet) served as positive and *PBS* (grey) as negative control. **(C, D)** *TEM* images of the functionalized DNA shells capturing multiple subviral particles **(C)** and *HBV* virions **(D)**. *HBV* particles were fixed with formaldehyde prior to incubation with DNA nanoshells. Scalebar indicates 100 nm.

**Figure 3.**
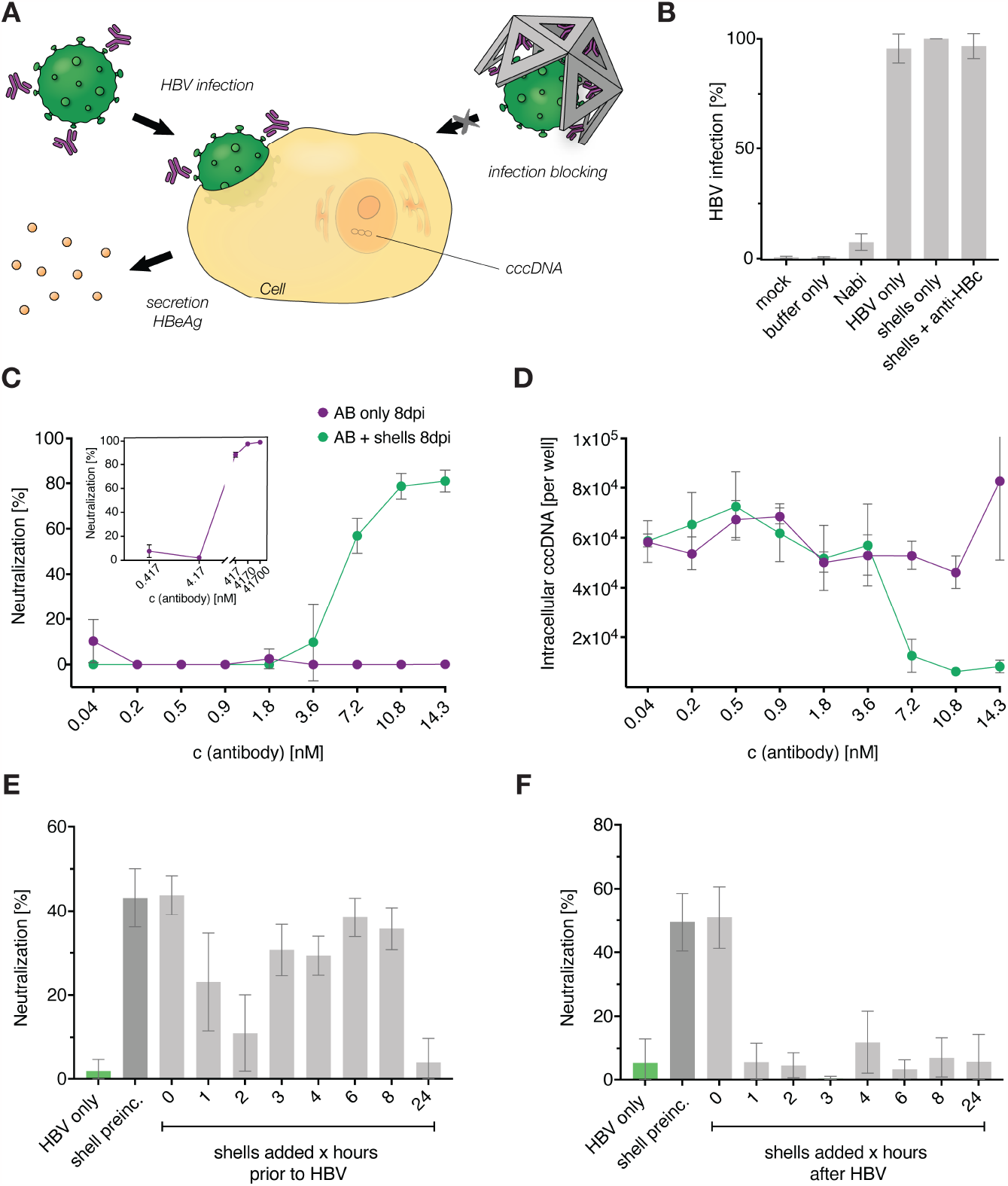
Hepatitis B virus neutralization. **(A)** Illustration of the neutralization assay comparing the neutralization capabilities of antibodies only versus DNA nanoshells functionalized with antibodies using wild-type *HBV*. If the host cell becomes infected, it starts secreting *HBeAg* (shown in yellow) and viral *cccDNA* can be found in the nucleus (indicated in orange). **(B-D)** *NTCP* expressing *HepG2* cells were infected with wild-type *HBV* and infection efficacy was analyzed 8 days after infection. **(B)** Infection efficacy determined by relative *HBeAg* secretion level quantified by *ELISA*. Background was determined using cell culture medium or the buffer used for nanoshell production. Infection neutralization by the therapeutically used human immune globulin *Nabi-HB*® is shown in comparison to the positive control of adding *HBV* only, *HBV* together with unfunctionalized nanoshells or nanoshells functionalized with polyclonal *anti-HBc* antibody, which should not bind *HBsAg*. Neutralization capacity is given in % based on *HBeAg* detection. **(C)** Neutralization capacity of nanoshells functionalized with *MoMAb* anti-*HBs antibody* (purple) compared to the free antibody (green). The insert shows the neutralization capacity of free antibody for a larger concentration range. Neutralization capacity is given in % based on *HBeAg* detection via *ELISA*. **(D)** Intracellular *HBV cccDNA* quantified by PCR. **(B-D)** The data is represented as mean ± *s*.*d*. of n = 3 independent experiments. **(E, F)** *HepG2-NTCP* cells were infected with r*HBV* expressing *Nano-luc* luciferase **(E)** Neutralization capacity of nanoshells administered to *HepG2-NTCP* cells at different time points before *HBV* infection. **(F)** Neutralization capacity of nanoshells administered to *HepG2-NTCP* cells at different time points after *HBV* infection. Shell preinc. indicates functionalized shells which were preincubated with HBV for 3h before administration to the cells. Infection efficacy and levels of neutralization were determined by luciferase activity in cell culture medium. Respective values determined in mock-treated cells served to define 0% neutralization. The data is represented as mean ± *s*.*d*. and is composed of n = 6 independent experiments.

To ensure stability under cell culture conditions, we coated the antibody-functionalized nanoshells with a mixture of polylysine and polyethylene-glycol-polylysine (*PEG-polylysine*)^27, 28^ a stabilization strategy that has previously been shown to stabilize similar DNA nanoshells for at least 24h.^12^ We confirmed the structural integrity using negative-staining transmission electron microscopy (*TEM*) (Figure 2A, Supplementary Figure 4A). To test the ability of the functionalized nanoshells to trap *HBV*, we added non-infectious subviral particles to the shells. We imaged them using *TEM*, which revealed dense packaging of the subviral particles inside the shell cavity (Figure 2C, Supplementary Figure 4B). We also attempted to trap purified infectious *HBV* virions using the shells. To this end, the *HBV* particles were fixed with paraformaldehyde and administered to the functionalized nanoshells. Due to the low concentration of the purified, enveloped *HBV* virions that we obtained compared to subviral particles or non-enveloped viruses, TEM imaging proved challenging (Figure 2D, Supplementary Figure 4C).

To determine the ability of the DNA nanoshells to neutralize infectious *HBV* in cell culture, we carried out experiments schematically depicted in Figure 3A. We quantified the degree of cellular infection by measuring secreted *HBeAg* levels and intracellular *cccDNA* using enzyme-coupled immunosorbent assays (*ELISA*) and quantitative *PCR* (*q-PCR*), respectively.

To normalize the data, we used the levels of these markers in control *HBV*-infection samples where *HBV* was allowed to infect and proliferate freely. The term “neutralization” denotes the fractional suppression of infection relative to those controls. We seeded *NTCP*-expressing *HepG2* cells in a differentiation medium 3 days before infecting them with *HBV* at a multiplicity of infection (*MOI*) of 100 enveloped, DNA-containing virus particles per cell. The infectious *HBV* particles were preincubated with either DNA nanoshells or free antibodies for 3 hours at 37 °C and then administered to the cells. After 18 hours, we removed the inoculum, washed the cells, and added fresh media. Cell culture supernatants were collected 4- and 8-days post-infection, respectively, to determine *HBeAg* and *cccDNA* levels.

To establish the robustness of the assay and assess the specificity of the functionalized nanoshells, we performed several control experiments. Mock and buffer samples not containing any *HBV* showed a 0% infection rate and were used to define a hypothetical 100% infection neutralization rate. Multiple biological replicates of *HBV*-only controls were used to set a 100% infection rate within the experimental variation. Non-functionalized nanoshells were incubated at equivalent stoichiometric ratios to *HBV* and did not interfere with infection, which was anticipated since nonfunctionalized shell variants should not interact with *HBV*. Finally, we prepared a functionalized version of the nanoshells coated with an *IgG* antibody recognizing *HBcAg* i.e., the capsid inside the virus particles that should not be accessible in the infectious and enveloped variant of *HBV*. Therefore, *anti-HBc IgG* functionalized nanoshells were expected to neither interact with the enveloped *HBV* nor to suppress infection. As expected, we observed negligible infection suppression using this control.

Next, we collected dose-response data for neutralization using both the *anti-HBs MoMAb* alone and the nanoshells functionalized with this antibody (Figure 3C, D). The data is presented as a function of the effective *MoMAb IgG* concentration to ensure a fair comparison. As 90 *IgG* molecules were coated per shell, the actual shell concentration is 90 times lower than indicated on the graphs. Our results show that the *MoMAb*-functionalized nanoshells potently neutralized the viruses with an estimated half maximal inhibitory concentration (*IC*_*50*_) of ∼5 nmol/L in terms of effective *IgG* antibody concentration, equivalent to an *IC*_*50*_ of ∼55 pmol/L in terms of actual DNA origami nanoshell concentration. When using free *MoMAb IgG* at 5 nmol/L instead of mounting them in groups of 90 on nanoshells, we observed negligible neutralization effects, with neutralization only occurring at ∼100-fold higher concentrations of around 0.5 µmol/L free *IgG* (Figure 3C inset). We also quantified the amount of intracellular *cccDNA* in the cells indicating the number of infected cells (Figure 3D). The amount of intracellular *cccDNA* dropped sharply in the dose-response curves obtained for the DNA nanoshells at an effective *IgG* concentration of ∼5 nM, which is consistent with our findings with the *HBeAg* assay (Figure 3C).

To investigate how a temporal offset between *HBV*-capturing shells and the addition of infectious *HBV* affects the neutralization capacity of the nanoshells, we infected cells with recombinant *HBV (rHBV)* expressing a secreted nano-luciferase. This allows more sensitive detection of *HBV* infection at a lower, more physiological *MOI. rHBV* was used at a *MOI* of 10 virions/cell and nanoshells were added at a concentration of 15.9 nmol/L at the time of infection or at different time offsets of 1, 2, 3, 4, 6, 8, and 24 hours before or after infection. To quantify infection, we measured luminescence in cell lysates 8 days post infection as a readout. We found that preincubation of shells with *HBV* and simultaneous addition of both components led to comparable levels of neutralization (Figure 3E) indicating that the nanoshells readily engulf the infectious virus in the cell culture medium even at high dilution.

The relative timing of addition of shells and viruses to cells impacted the neutralization capacity of the nanoshells. If shells were added prior to the virus, the degree of neutralization decreased with increasing temporal offset (Figure 3F). This was most likely due to progressive degradation of the shell material in the cell culture environment.^12^ On the other hand, administering shells a posteriori to exposing cells to viruses did not neutralize viruses for temporal offsets of 1 hour or longer. This finding makes sense considering that the closed cell contact by *HBV* is expected to be established between 30 to 60 minutes.^29^ Once the viruses have entered the cells, they can no longer be sequestered by the *HBV*-capturing shells as these shells are not expected to enter the cell.

## Conclusions

In previous work,^12^ we developed design principles and methods to construct DNA origami nanoshells and successfully trapped various viruses using different types of internal functionalization.^13 30^ In the present study we show the first successful in vitro neutralization of a live human pathogen using these shells, whereas in previous work non-infectious model particles such as the HBV core and non-replicative viruses were used. An additional distinctive feature is the achievement of neutralization enhancement of the efficacy of the recombinant *IgG* antibody *MoMAb*, which we used as a coating inside the shells to specifically trap *HBV*, significantly from an *IC*_*50*_ of around 0.5 µmol/L^25^ per free *anti-HBs IgG* to ∼5 nmol/L when the *IgG* is mounted inside the shells, corresponding to an *IC*_*50*_ of approximately 55 pmol/L of *HBV*-capturing shells. Consequently, our approach thus allows for a two orders of magnitude reduction in the required antibody quantity. The potency enhancements are presumably achieved by two mechanisms working in concert: multivalent binding of *HBV* virions to the *anti-HBs* antibody inside shells, which leads to avidity effects, and steric occlusion by the shell material, creating a physical barrier that prevents the virus from interacting with host cells. The latter was confirmed by a clear temporal offset when nanoshells were added at distinct time points before or after the virus. The medications available for HBV, particularly those that target the HBsAg, are often sidetracked by the substantial secretion of SVPs from infected cells^20^. This is surely also the case for DNA nanoshells; however, as the TEM images suggest (Figure 2B), each nanoshell has the capacity to incorporate a substantial number of SVPs. This could potentially enhance their efficacy by reducing the number of SVPs in solution and allowing actual hepatitis virions to be captured.

Our findings establish the shell system as a molecular framework for constructing new antivirals from components that have weak antiviral properties when used individually. Whether the nanoshell-virus complexes are taken up by antigen-presenting cells and inducing a virus-specific T-cell response, remains to be investigated. First results hint toward the uptake of the complexes by monocyte derived dendritic cells (moDC) (Supplementary Figure 7) but need further approvement. Another proposed advantage of our shells is the fact that they should be less prone to antibody dependent infection enhancement, a mechanism where antibodies are not able to efficiently eliminate infected cells/viruses but where the antibody-virus complex is taken up by the cells, so the virus is able to replicate within the cell. *In vivo* efficacy studies are crucial next steps to establish the therapeutic potential of the DNA shell system.

## Materials & Methods

### DNA origami shell design, self-assembly, and purification

The triangular shell subunits were designed and self-assembled as previously described^12^ using a scaffold with a length of 8064 bases.^31^ The resulting triangular monomers were purified using agarose gel purification with an agarose concentration of 1.5% and a 0.5x*TBE* buffer containing 5.5 mM *MgCl*_*2*_. The leading bands were extracted from the gel and the agarose removed using the Corning® Costar® Spin-X® centrifuge tube filters with a cellulose acetate membrane and a pore size of 0.45 μm. Residual agarose was removed by spinning the sample for 35 min at 21000 rcf and collecting the supernatant. Details of the purification methods can be found in.^32^ The purified monomers were assembled to T=1 nanoshells by increasing the *MgCl*_*2*_ concentration in the buffer to 40 mM and incubating at 40 °C for at least 8 days. The resulting T=1 nanoshells are made up of 10 triangular subunits, each with 9 single stranded DNA sequences (“handles”) protruding into the cavity of the shell.

### Shell stabilization

The T=1 nanoshells were stabilized as described previously.^12^ This includes the strategy of UV point welding^26^ the individual subunit protrusions and recesses to ensure the structural integrity of the shell. The UV light source used is a custom build UV lamp with the deep ultraviolet light emission source (*DUVLED*) *DUV310-SD353EN* from Roither Laser Technik GmbH. The *DUVLED* emits light at a typical peak wavelength of 310 nm and is operated at an optical output power of 43 mW at 350 mA. For the irradiation procedure the sample is placed into self-made Teflon cuvette to maximize the reflection of the incoming UV light. All T=1 nanoshell samples were irradiated 10 min with UV light prior to functionalization with antibodies. After functionalization the construct was further stabilized by coating the structure using a 0.6:1 *N/P* ratio with a 1:1 mixture of *K*_*10*_-oligolysine (*PL*) and *K*_*10*_-*PEG*_*5K*_-oligolysine (*PPL*), a stabilization strategy described perviously^27^ and modified to stabilize higher order DNA origami structures.

### DNA coupling of anti-HBs antibodies

The *anti-S scFV-Linker-hIgG1Fcmut* antibodies (*MoMAb*) used to capture subviral particles and *HBV* virions were previously described and characterized^25^. Large scale production was contracted to InVivo Biotech Services GmbH. The produced antibodies were modified with a 26 bp DNA sequence, complementary to the handle sequence inside the nanoshells:

**Table 1.**
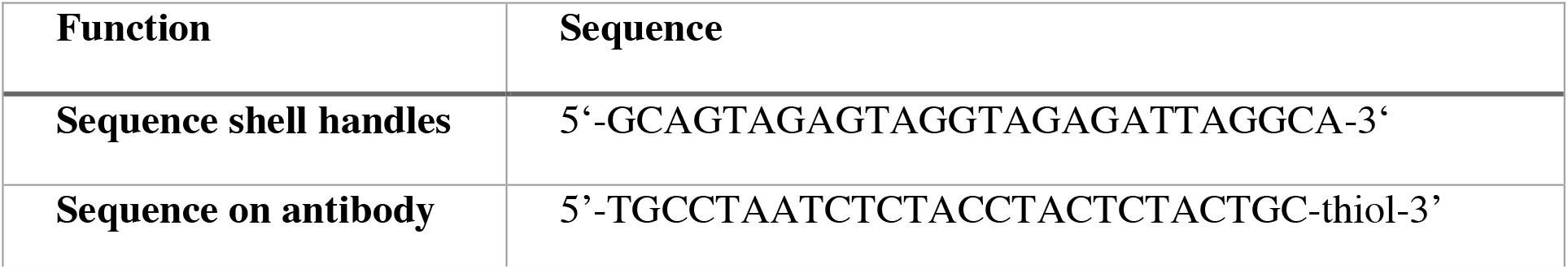
DNA sequences for antibody attachment

The modification was accomplished by connecting the thiol modified DNA strand to the antibody via a *Sulfo-SMCC* crosslinker from Thermo Scientific™, using a ratio of antibody to DNA strand of 1:7 (Supplementary Figure 2, 3). The resulting mixture was purified using the ion-exchange chromatography system proFire by Dynamic Biosensors.

### Nanoshells and *HBV* subviral particles and virion binding

The nanoshells were incubated with the antibodies in a handle to antibody ratio of 1:1 and incubated overnight at 25 °C. They were subsequently coated with the *PL/PPL* coating for at least 2 h. For *TEM* imaging we added subviral particles in *PBS* buffer or purified Hepatitis B viruses to the functionalized nanoshells and incubated for at least 4 h. For neutralization experiments the functionalized nanoshell samples or the antibody only samples were incubated with the Hepatitis B virus sample at different ratios at 37 °C for 3 h.

### Negative staining *TEM*

For *TEM* imaging the samples were incubated on *FCF-400-Cu* TEM grids from Electron Microscopy Sciences with a Formvar Carbon Film, which was previously glow-discharged for 45 s with a charge of 35 mA. The incubation time ranged from 1 min to 20 min depending on the concentration of the sample. The samples were subsequently submerged in 2 % aqueous uranyl formate solution containing 25 mM sodium hydroxide for staining and blotted dry with filter paper. Images were acquired using a FEI Tecnai T12 microscope at 120 kV and a Tietz TEMCAM-F416 camera, all operated with the Software SerialEM. The magnification of the images ranged from 21000x to 52000x. The contrast of the images was subsequently globally enhanced with Fiji ^33^ to show details of individual features.

### Neutralization assays

To determine the neutralization capacity of the nanoshells, *HepG2-NTCP* cells were seeded in a collagenized 24-well plate at a density of 3·10^5^ cells/well in differentiation medium, which is *DMEM* high glucose supplemented with 10% *FCS*, 2.5% *DMSO, Pen/Strep*, non-essential amino acids, L-glutamine, and sodium pyruvate (all Gibco Life Technologies) 3 days before infection. Different concentrations of nanoshells or *MoMAb* antibodies were mixed with purified *HBV*,^*34*^ genotype D and incubated at 37 °C for 3 hours. *PEG* was added to reach a final concentration of 4% (v/v) before adding the virus-nanoshell mixture to the cells at an *MOI* of 100 viral particles per cell. After 18 hours, the inoculum was removed, cells were washed twice with *PBS*, and 1 mL of medium was added to the cells for further cultivation. At days 4 and 8 post infection, cell culture medium was collected, and *HBeAg* levels were determined by *ELISA* to calculate the neutralization capacity. In addition, cells were lysed 8 days post infection to determine intracellular *HBV cccDNA* and *rcDNA* via *qPCR*.

To determine the prophylactic potential of the nanoshells, shells were added at different time points before infection of the *HepG2-NTCP* cells with a luciferase-expressing *rHBV* (genotype D, *MOI* 10 viral particles/cell) in the presence of 4 % *PEG* (v/v). To determine the therapeutic potential, nanoshells were added at different time points after infection. 18 hours after infection, the inoculum was removed, the cells were washed twice with *PBS*, and 1 mL media was added to the cells. At days 4 and 8 post infection, luciferase activity of 100 µL cell culture medium was determined to calculate neutralization capacity.

## Supporting Information

Additional information on size comparisons, DNA-antibody coupling, overviews of negative stain *EM* data, dose response curves, and uptake by dendritic cells is given in the supporting information.

## Author Contributions

E.M.W and F.K. contributed equally to this work. H.D. and U.P. designed and co-supervised the research. E.M.W. performed shell production, stabilization, antibody conjugation and the final shell modification. F.M. designed *MoMAb*. E.M.W. and F.K. performed control experiments to determine *MoMAb* functionality and successful virus binding. F.K. and U.P. provided *MoMAb*, subviral particles and hepatitis B virions. F.K. performed cell culture neutralization experiments. F.K. and E.M.W. analyzed data.

## Acknowledgements

This work has received funding from the European Union’s Horizon 2020 research and innovation program within the *FET* Open project Virofight (grant agreement No 899619, to HD). This work was further supported by the Deutsche Forschungsgemeinschaft via the Gottfried-Wilhelm-Leibniz Program (to HD), via grant ID DI1500/5 (to HD) and via CRC-TRR179 (project No. 272983813, to UP). We thank C. Sigl, J. Kretzmann and T. Gerling for technical discussions.

