## Supplemental Information for "Hepatitis B virus neutralization with DNA origami nanoshells"

**Content**

Supplementary Figure 1: 3D rendering of the *HBV* and the T=1 nano-shell

Supplementary Figure 2: DNA-antibody coupling with *Sulfo-SMCC*

Supplementary Figure 3: Negative stain electron microscopy images

Supplementary Figure 4: Agarose gel of the purified DNA-antibody product

Supplementary Figure 5: Dose-response of *AB* only vs T=1 nano-shell neutralization

Supplementary Figure 6: Statistically significant neutralization

Supplementary Figure 7: Nanoshell uptake by dendritic cells

Supplementary Note 1: Scaffold sequence.

Supplementary Note 2: Staple sequences triangular monomer 1

Supplementary Note 3: Staple sequences triangular monomer 2

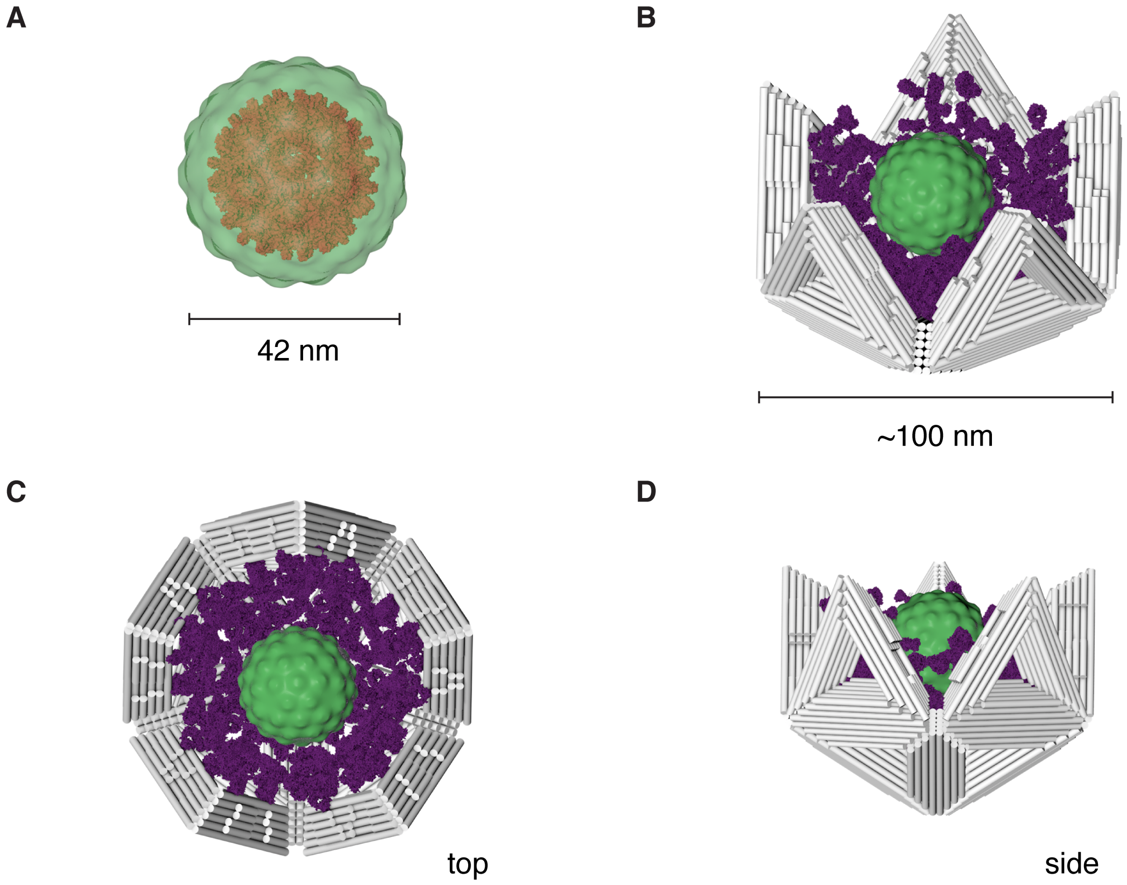


**Supplementary figure 1**. **3D rendering of the *HBV* and the T=1 nano-shell.** 3D rendering of **(A)** the Hepatitis B virus with its core particle colored in red and the envelope colored in green. To scale rendering of the **(B)** Hepatitis B virus embedded in the T=1 nano-shell. **(C)** View from the top and **(D)** view from the side.

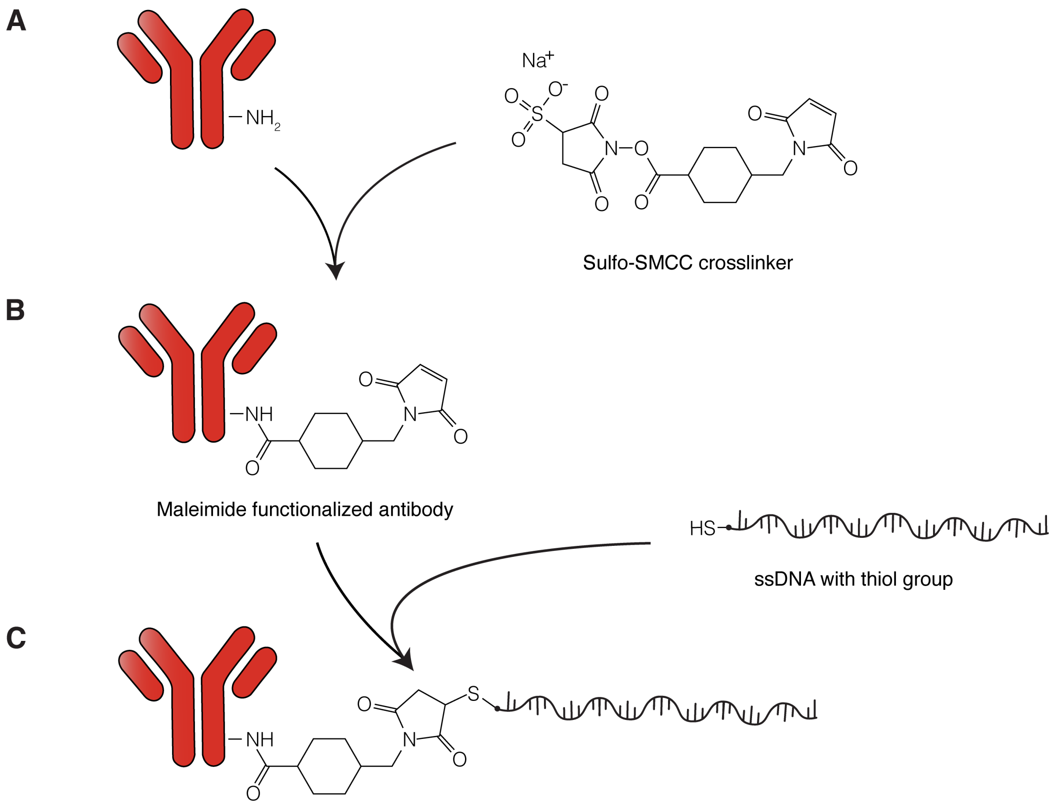


**Supplementary figure 2**. **Illustration of the DNA-antibody coupling with *Sulfo-SMCC*.**

**(A)** Antibody with free amine groups is mixed with a *Sulfo-SMCC* crosslinker. **(B)** Maleimide functionalized antibody is incubated with a thiolated single-stranded DNA handle. **(C)** Final DNA functionalized antibody product. Adapted from [1].

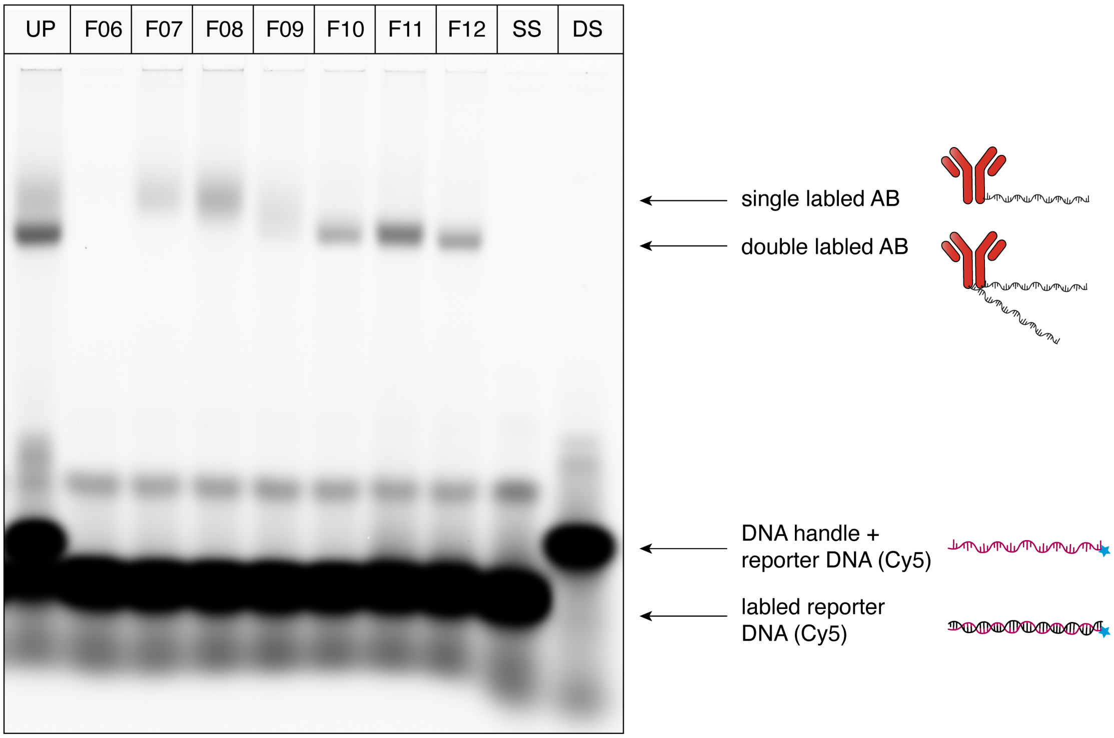


**Supplementary figure 3**. **Agarose gel of the purified DNA-antibody product.** 4% agarose gel analysis of the DNA-antibody product after purification with ion-exchange chromatography. *UP* stands for the unpurified product. *F06 – F12* stands for the purification fractions. *SS* is a single stranded DNA reporter with Cyanine 5 dye, complementary to the handle on the antibody. *DS* is the *SS* reporter with the complementary handle without the antibody.

A

B

C

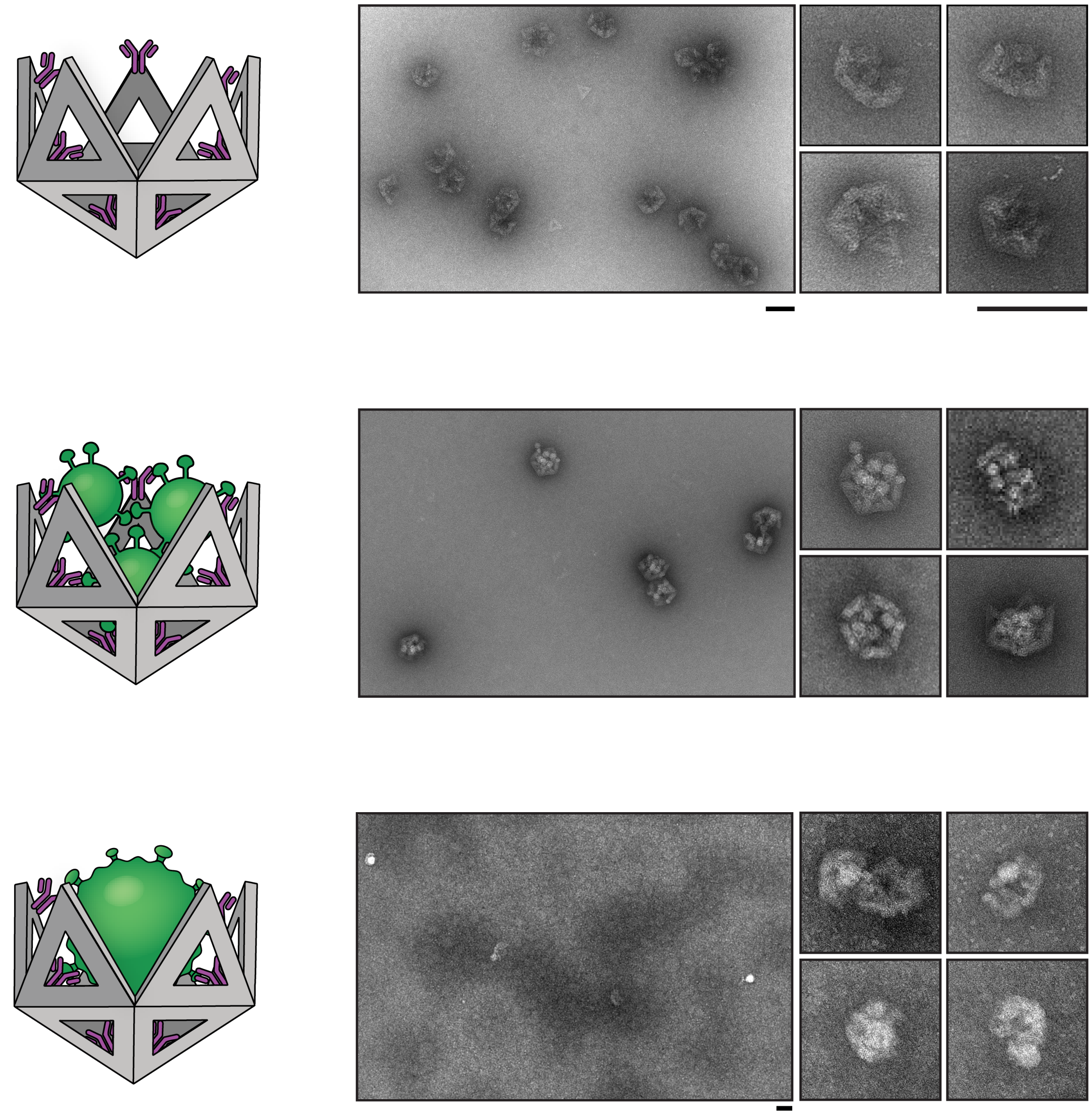

**Supplementary figure 4**. **Negative stain electron microscopy images.** Overview of the **(A)** antibody functionalized DNA nano-shells, **(B)** the capture of Hepatitis B subviral particles and **(C)** the capture of *HBV* viruses fixed with formaldehyde. Scale bar indicates 100 nm.

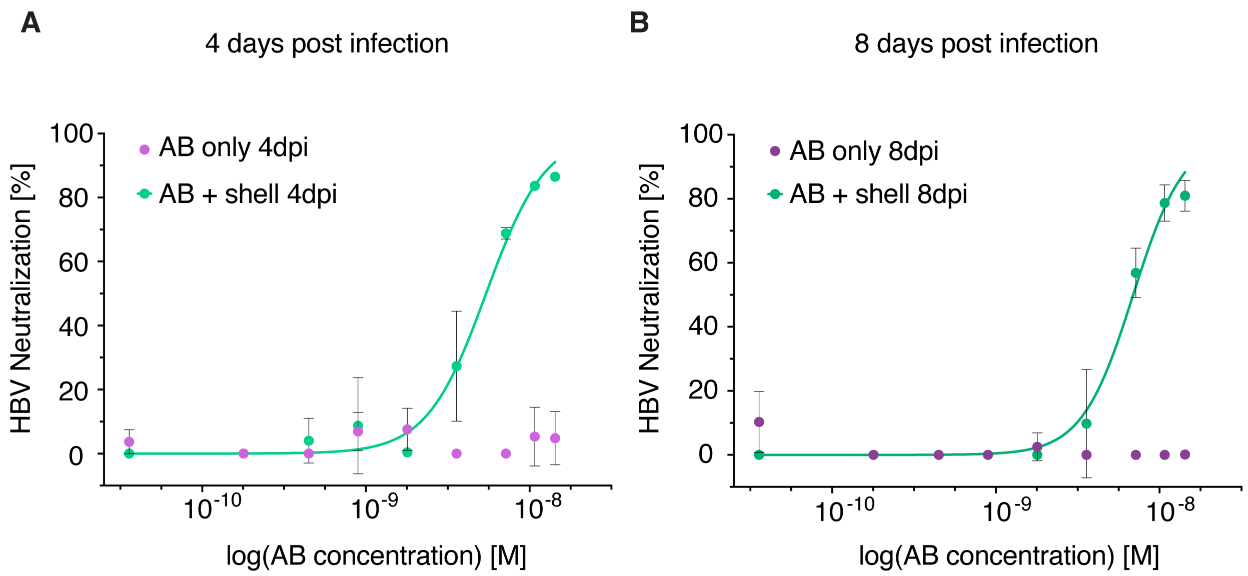


**Supplementary figure 5**. **Dose-response of *AB* only vs T=1 nano-shell neutralization**. *IC50* curves of the *HBV* neutralization data using functionalized nano-shells using the *HBeAg* detection. **(A)** Shows the neutralization capacity 4 days after the *HBV* infection (*4dpi*) and **(B)** 8 days after infection (*8dpi*). Neutralization data was fitted using a nonlinear regression and resulted in IC50 values of 5.4 nM ± 1.3 nM and 6.9 nM ± 1.2 nM for *4dpi* and *8dpi* respectively. The data is represented as mean + s.d. and is composed of n=3 biologically independent experiments.

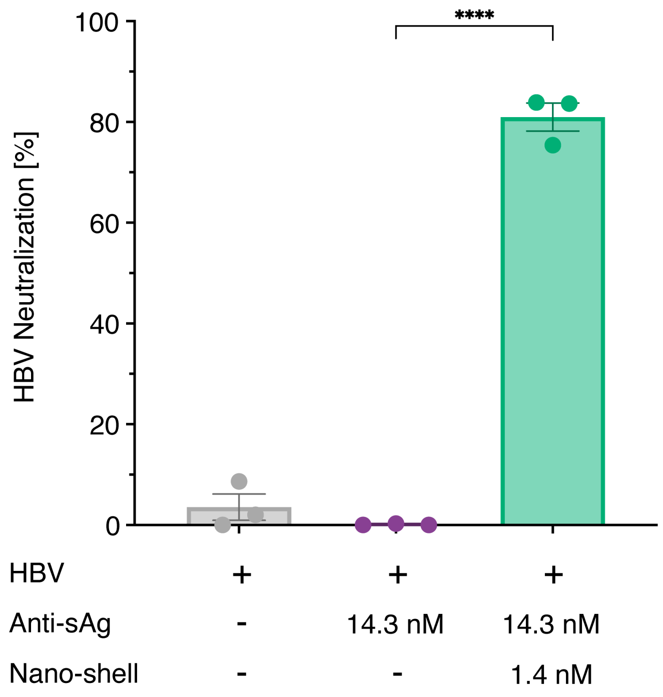


**Supplementary figure 6**. **Statistically significant neutralization.** One-way analysis of variance (*ANOVA*) of the highest tested nano-shell concentration to study statistically significant virus blocking compared with the antibody only sample.The functionalized nano-shells show a statistically significant higher neutralization capacity than the sample using the antibodies only (**** $P < 0.0001$).

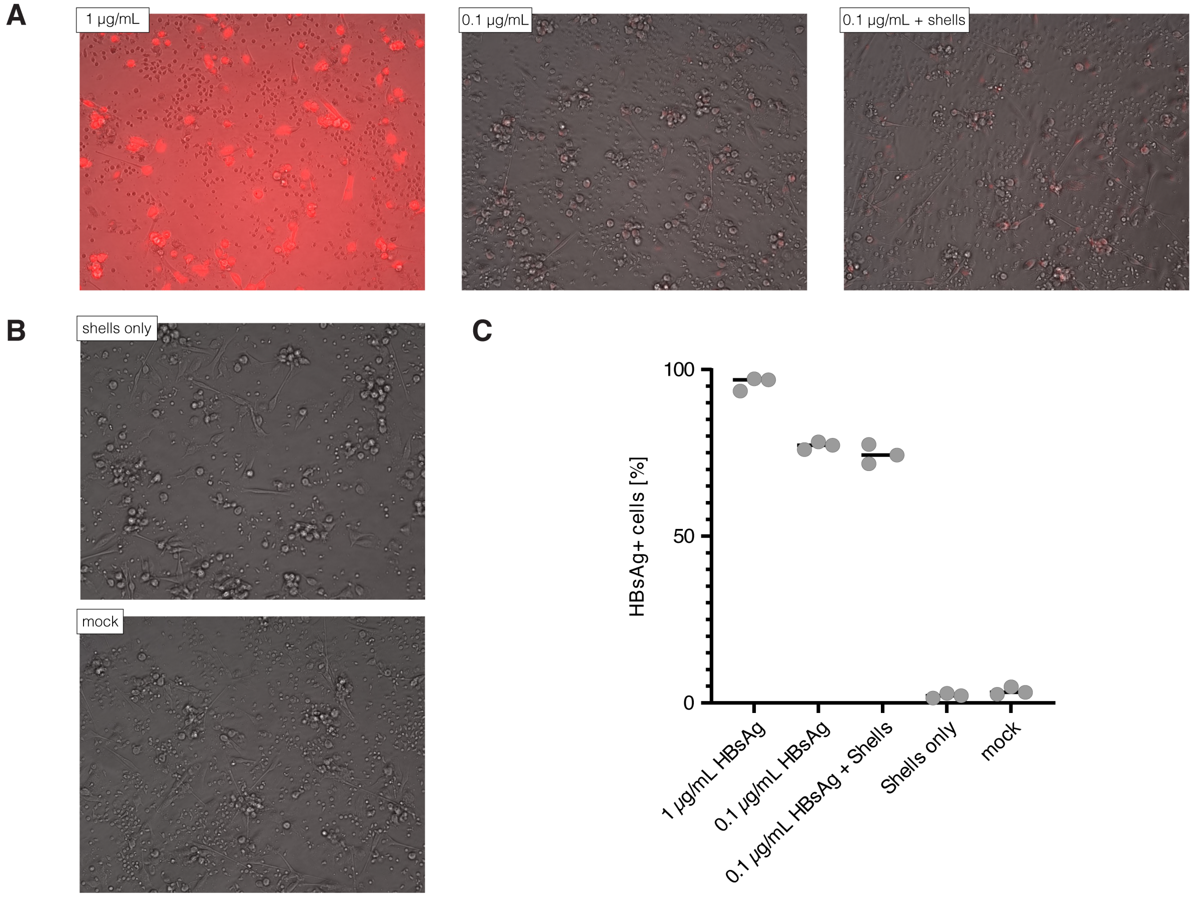


**Supplementary figure 7. Nano-shell uptake by dendritic cells.** (**A**) Nano-shells with fluorescently labeled *HBsAg* at different concentrations added to dendritic cells (*moDC*) **(B)** Cell control with shells only and mock. **(C)** Data on the amount of internalized HBsAg into the *moDC* cell line using n = 3 biological independent measurements.

**Supplementary Note 1 | Scaffold sequence**.

*M13MP18* modified sequence with length of 8064 bases.[2]

GGCAATGACCTGATAGCCTTTGTAGATCTCTCAAAAATAGCTACCCTCTCCGGCATT
AATTTATCAGCTAGAACGGTTGAATATCATATTGATGGTGATTTGACTGTCTCCGGC
CTTTCTCACCCTTTTGAATCTTTACCTACACATTACTCAGGCATTGCATTTAAAATAT
ATGAGGGTTCTAAAAATTTTTATCCTTGCGTTGAAATAAAGGCTTCTCCCGCAAAAG
TATTACAGGGTCATAATGTTTTTGGTACAACCGATTTAGCTTTATGCTCTGAGGCTTT
ATTGCTTAATTTTGCTAATTCTTTGCCTTGCCTGTATGATTTATTGGATGTTAATGCTA
CTACTATTAGTAGAATTGATGCCACCTTTTCAGCTCGCGCCCCAAATGAAAATATAG
CTAAACAGGTTATTGACCATTTGCGAAATGTATCTAATGGTCAAACTAAATCTACTC
GTTCGCAGAATTGGGAATCAACTGTTATATGGAATGAAACTTCCAGACACCGTACTT
TAGTTGCATATTTAAAACATGTTGAGCTACAGCATTATATTCAGCAATTAAGCTCTA
AGCCATCCGCAAAAATGACCTCTTATCAAAAGGAGCAATTAAAGGTACTCTCTAATC
CTGACCTGTTGGAGTTTGCTTCCGGTCTGGTTCGCTTTGAAGCTCGAATTAAAACGC
GATATTTGAAGTCTTTCGGGCTTCCTCTTAATCTTTTTGATGCAATCCGCTTTGCTTCT
GACTATAATAGTCAGGGTAAAGACCTGATTTTTGATTTATGGTCATTCTCGTTTTCTG
AACTGTTTAAAGCATTTGAGGGGGATTCAATGAATATTTATGACGATTCCGCAGTAT
TGGACGCTATCCAGTCTAAACATTTTACTATTACCCCCTCTGGCAAAACTTCTTTTGC
AAAAGCCTCTCGCTATTTTGGTTTTTATCGTCGTCTGGTAAACGAGGGTTATGATAGT
GTTGCTCTTACTATGCCTCGTAATTCCTTTTGGCGTTATGTATCTGCATTAGTTGAAT
GTGGTATTCCTAAATCTCAACTGATGAATCTTTCTACCTGTAATAATGTTGTTCCGTT
AGTTCGTTTTATTAACGTAGATTTTTCTTCCCAACGTCCTGACTGGTATAATGAGCCA
GTTCTTAAAATCGCATAAGGTAATTCACAATGATTAAAGTTGAAATTAAACCATCTC
AAGCCCAATTTACTACTCGTTCTGGTGTTTCTCGTCAGGGCAAGCCTTATTCACTGAA
TGAGCAGCTTTGTTACGTTGATTTGGGTAATGAATATCCGGTTCTTGTCAGATTACTC
TTGATGAAGGTCAGCCAGCCTATGCGCCTGGTCTGTACACCGTTCACTGTCCTCTTTC
AAAGTTGGTCAGTTCGGTTCCCTTATGATTGACCGTCTGCGCCTCGTTCCGGCTAAGT
AACATGGAGCAGGTCGCGGATTTCGACACAATTTATCAGGCGATGATACAAATCTCC
GTTGTACTTTGTTTCGCGCTTGGTATAATCGCTGGGGGTCAAAGATGAGTGTTTTAGT
GTATTCTTTTGCCTCTTTCGTTTTAGGTTGGTGCCTTCGTAGTGGCATTACGTATTTTA
CCCGTTTAATGGAAACTTCCTCATGAAAAAGTCTTTAGTCCTCAAAGCCTCTGTAGC
CGTTGCTACCCTCGTTCCGATGCTGTCTTTCGCTGCTGAGGGTGACGATCCCGCAAA
AGCGGCCTTTAACTCCCTGCAAGCCTCAGCGACCGAATATATCGGTTATGCGTGGGC
GATGGTTGTTGTCATTGTGGCGCAACTATCGGTATCAAGCTGTTTAAGAAATTCACC
TCGAAAGCAAGCTGATAAACCGATACAATTAAAGGCTCCTTTTGGAGCCTTTTTTTT
GGAGATTTTCAACGTGAAAAAATTATTATTCGCAATTCCTTTAGTTGTTCCTTTCTAT
TCTCACTCCGCTGAAACTGTTGAAAGTTGTTTAGCAAAATCCCATACAGAAAATTCA
TTTACTAACGTCTGGAAAGACGACAAAACTTTAGATCGTTACGCTAACTATGAGGGC
TGTCTGTGGAATGCTACAGGCGTTGTAGTTTGTACTGGTGACGAAACTCAGTGTTAC
GGTACATGGGTTCCTATTGGGCTTGCTATCCCTGAAAATGAGGGTGGTGGCTCTGAG
GGTGGCGGTTCTGAGGGTGGCGGTTCTGAGGGTGGCGGTACTAAACCTCCTGAGTAC
GGTGATACACCTATTCCGGGCTATACTTATATCAACCCTCTCGACGGCACTTATCCG
CCTGGTACTGAGCAAAACCCCGCTAATCCTAATCCTTCTCTTGAGGAGTCTCAGCCT
CTTAATACTTTCATGTTTCAGAATAATAGGTTCCGAAATAGGCAGGGGGCATTAACT
GTTTATACGGGCACTGTTACTCAAGGCACTGACCCCGTTAAAACTTATTACCAGTAC

ACTCCTGTATCATCAAAAGCCATGTATGACGCTTACTGGAACGGTAAATTCAGAGAC
TGCGCTTTCCATTCTGGCTTTAATGAGGATTTATTTGTTTGTGAATATCAAGGCCAAT
CGTCTGACCTGCCTCAACCTCCTGTCAATGCTGGCGGCGGCTCTGGTGGTGGTTCTG
GTGGCGGCTCTGAGGGTGGTGGCTCTGAGGGTGGCGGTTCTGAGGGTGGCGGCTCT
GAGGGAGGCGGTTCCGGTGGTGGCTCTGGTTCCGGTGATTTTGATTATGAAAAGATG
GCAAACGCTAATAAGGGGGCTATGACCGAAAATGCCGATGAAAACGCGCTACAGTC
TGACGCTAAAGGCAAACTTGATTCTGTCGCTACTGATTACGGTGCTGCTATCGATGG
TTTCATTGGTGAGTTTCCGGCCTTGCTAATGGTAATGGTGCTACTGGTGATTTTGCTG
GCTCTAATTCCCAAATGGCTCAAGTCGGTGACGGTGATAATTCACCTTTAATGAATA
ATTTCCGTCAATATTTACCTTCCCTCCCTCAATCGGTTGAATGTCGCCCTTTTGTCTTT
GGCGCTGGTAAACCATATGAATTTTCTATTGATTGTGACAAAATAAACTTATTCCGT
GTGTCTTTGCGTTTCTTTTATATGTTGCCACCTTTATGTATGTATTTTCTACGTTTGCT
AACATACTGCGTAATAAGGAGTCTTAATCATGCCAGTTCTTTTGGGTATTCCGTTATT
ATTGCGTTTCCTCGGTTTCCTTCTGGTAACTTTGTTCGGCTATCTGCTTATTTTCTTAA
AAAGGGCTTCGGTAAGATAGCTATTGCTATTTCATTGTTTCTTGCTCTTATTATTGGG
CTTAACTCAATTCTTGTGGGTTATCTCTCTGATATTAGCGCTCAATACCCTCTGACTT
TGTTCAGGGTGTTCAGTTAATTCTCCCGTCTAATGCGCTTCCCTGTTTTTATGTTATTC
TCTCTGTAAAGGCTGCTATTTTCATTTTTGACGTTAAACAAAAAATCGTTTCTTATTT
GGATTGGGATAAATAATATGGCTGTTTATTTTGTAACTGGCAAATTAGGCTCTGGAA
AGACGCTCGTTAGCGTTGGTAAGATTCAGGATAAAATTGTAGCTGGGTGCAAAATA
GCAACTAATCTTGATTTAAGGCTTCAAAACCTCCCGCAAGTCGGGAGGTTCGCTAAA
ACGCCTCGCGTTCTTAGAATACCGGATAAGCCTTCTATATCTGATTTGCTTGCTATTG
GGCGCGGTAATGATTCCTACGATGAAAATAAAAACGGCTTGCTTGTTCTCGATGAGT
GCGGTACTTGGTTTAATACCCGTTCTTGGAATGATAAGGAAAGACAGCCGATTATTG
ATTGGTTTCTACATGCTCGTAATTAGGATGGGATATTATTTTTCTTGTTCAGGACTTA
TCTATTGTTGATAAACAGGCGCGTTCTGCATTAGCTGAACATGTTGTTTATTGTCGTC
GTCTGGACAGAATTACTTTACCTTTTGTCGGTACTTTATATTCTCTTATTACTGGCTC
GAAAATGCCTCTGCCTAAATTACATGTTGGCGTTGTTAAATATGGCGATTCTCAATT
AAGCCCTACTGTTGAGCGTTGGCTTTATACTGGTAAGAATTTGTATAACGCATATGA
TACTAAAAGGCTTTTTCTAGTAATTATGATTCCGGTGTTTATTCTTATTTAACGCCTT
ATTTATCACACGGTCGGTATTTCAAACCATTAAATTTAGGTCAGAAGATGAAATTAA
CTAAAATATATTTGAAAAAGTTTTCTCGCGTTCTTTGTCTTGCGATTGGATTTGCATC
GCATTTACATATAGTTATATAACCCAACCTAAGCCGGAGGTTAAAAAGGTAGTCTCT
CAGACCTATGATTTTGATAAATTCACTATTGACTCTTCTCAGCGTCTTAATCTAAGCT
ATCGCTATGTTTTCAAGGATTCTAAGGGAAAATTAATTAATAGCGACGATTTACAGA
AGCAAGGTTATTCACTCACATATATTGATTTATGTACTGTTTCCATTAAAAAAGGTA
ATTCAAATGAAATTGTTAAATGTAATTAATTTTGTTTTCTTGATGTTTGTTTCATCATC
TTCTTTTGCTCAGGTAATTGAAATGAATAATTCGCCTCTGCGCGATTTTGTAACTTGG
TATTCAAAGCAATCAGGCGAATCCGTTATTGTTTCTCCCGATGTAAAAGGTACTGTT
ACTGTATATTCATCTGACGTTAAACCTGAAAATCTACGCAATTTCTTTATTTCTGTTT
TACGTGCAAATAATTTTGATATGGTAGGTTCTAACCCTTCCATTATTCAGAAGTATA
ATCCAAACAATCAGGATTATATTGATGAATTGCCATCATCTGATAATCAGGAATATG
ATGATAATTCCGCTCCTTCTGGTGGTTTCTTTGTTCCGCAAAATGATAATGTTACTCA
AACTTTTAAAATTAATAACGTTCGGGCAAAGGATTTAATACGAGTTGTCGAATTGTT
TGTAAAGTCTAATACTTCTAAATCCTCAAATGTATTATCTATTGACGGCTCTAATCTA
TTAGTTGTTAGTGCTCCTAAAGATATTTTAGATAACCTTCCTCAATTCCTTTCAACTG

TTGATTTGCCAACTGACCAGATATTGATTGAGGGTTTGATATTTGAGGTTCAGCAAG
GTGATGCTTTAGATTTTTCATTTGCTGCTGGCTCTCAGCGTGGCACTGTTGCAGGCGG
TGTTAATACTGACCGCCTCACCTCTGTTTTATCTTCTGCTGGTGGTTCGTTCGGTATTT
TTAATGGCGATGTTTTAGGGCTATCAGTTCGCGCATTAAAGACTAATAGCCATTCAA
AAATATTGTCTGTGCCACGTATTCTTACGCTTTCAGGTCAGAAGGGTTCTATCTCTGT
TGGCCAGAATGTCCCTTTTATTACTGGTCGTGTGACTGGTGAATCTGCCAATGTAAA
TAATCCATTTCAGACGATTGAGCGTCAAAATGTAGGTATTTCCATGAGCGTTTTTCCT
GTTGCAATGGCTGGCGGTAATATTGTTCTGGATATTACCAGCAAGGCCGATAGTTTG
AGTTCTTCTACTCAGGCAAGTGATGTTATTACTAATCAAAGAAGTATTGCTACAACG
GTTAATTTGCGTGATGGACAGACTCTTTTACTCGGTGGCCTCACTGATTATAAAAAC
ACTTCTCAGGATTCTGGCGTACCGTTCCTGTCTAAAATCCCTTTAATCGGCCTCCTGT
TTAGCTCCCGCTCTGATTCTAACGAGGAAAGCACGTTATACGTGCTCGTCAAAGCAA
CCATAGTACGCGCCCTGTAGCGGCGCATTAAGCGCGGCGGGTGTGGTGGTTACGCG
CAGCGTGACCGCTACACTTGCCAGCGCCCTAGCGCCCGCTCCTTTCGCTTTCTTCCCT
TCCTTTCTCGCCACGTTCGCCGGCTTTCCCCGCAAGCTCTAAATCGGGGGCTCCCTTT
AGGGTTCCGATTTAGTGCTTTACGGCACCTCGACCCCAAAAAACTTGATTTGGGTGA
TGGTTCACGTAGTGGGCCATCGCCCTGATAGACGGTTTTTCGCCCTTTGACGTTGGA
GTCCACGTTCTTTAATAGTGGACTCTTGTTCCAAACTGGAACAACACTCAACCCTAT
CTCGGGCTATTCTTTTGATTTATAAGGGATTTTGCCGATTTCGGAACCACCATCAAAC
AGGATTTTCGCCTGCTGGGGCAAACCAGCGTGGACCGCTTGCTGCAACTCTCTCAGG
GCCAGGCGGTGAAGGGCAATCAGCTGTTGCCCGTCTCACTGGTGAAAAGAAAAACC
ACCCTGGCGCCCAATACGCAAACCGCCTCTCCCCGCGCGTTGGCCGATTCATTAATG
CAGCTGGCACGACAGGTTTCCCGACTGGAAAGCGGGCAGTGAGCGCAACGCAATTA
ATGTGAGTTAGCTCACTCATTAGGCACCCCAGGCTTTACACTTTATGCTTCCGGCTCG
TATGTTGTGTGGAATTGTGAGCGGATAACAATTTCACACAGGAAACAGCTATGACCA
TGATTACGAATTCGAGCTCGGTACCCGGGGATCCTCAACTGTGAGGAGGCTCACGG
ACGCGAAGAACAGGCACGCGTGCTGGCAGAAACCCCCGGTATGACCGTGAAAACG
GCCCGCCGCATTCTGGCCGCAGCACCACAGAGTGCACAGGCGCGCAGTGACACTGC
GCTGGATCGTCTGATGCAGGGGGCACCGGCACGCTGGCTGCAGGTAACCCGGCATC
TGATGCCGTTAACGATTTGCTGAACACACCAGTGTAAGGGATGTTTATGACGAGCAA
AGAAACCTTTACCCATTACCAGCCGCGGGCAACAGTGACCCGGCTCATACCGCAAC
CGCGCCCGGCGGATTGAGTGCGAAAGCGCCTGCAATGACCCCGCTGATGCTGGACA
CCTCCAGCCGTAAGCTGGTTGCGTGGGATGGCACCACCGACGGTGCTGCCGTTGGCA
TTCTTGCGGTTGCTGCTGACCAGACCAGCACCACGCTGACGTTCTACAAGTCCGGCA
CGTTCCGTTATGAGGATGTGCTCTGGCCGGAGGCTGCCAGCGACGAGACGAAAAAA
CGGACCGCGTTTGCCGGAACGGCAATCAGCATCGTTTAACTTTACCCTTCATCACTA
AAGGCCGCCTGTGCGGCTTTTTTTACGGGATTTTTTTATGTCGATGTACACAACCGCC
CAACTGCTGGCGGCAAATGAGCAGAAATTTAAGTTTGATCCGCTGTTTCTGCGTCTC
TTTTTCCGTGAGAGCTATCCCTTCACCACGGAGAAAGTCTATCTCTCACAAATTCCG
GGACTGGTAAACATGGCGCTGTACGTTTCGCCGATTGTTTCCGGTGGGTTATCCGTT
CCCGTGGCGGCTCCACCTCTGAAAGCTTGGCACTGGCCGTCGTTTTACAACGTCGTG
ACTGGGAAAACCCTGGCGTTACCCAACTTAATCGCCTTGCAGCACATCCCCCTTTCG
CCAGCTGGCGTAATAGCGAAGAGGCCCGCACCGATCGCCCTTCCCAACAGTTGCGC
AGCCTGAATGGCGAATGGCGCTTTGCCTGGTTTCCGGCACCAGAAGCGGTGCCGGA
AAGCTGGCTGGAGTGCGATCTTCCTGAGGCCGATACTGTCGTCGTCCCCTCAAACTG
GCAGATGCACGGTTACGATGCGCCCATCTACACCAACGTGACCTATCCCATTACGGT

CAATCCGCCGTTTGTTCCCACGGAGAATCCGACGGGTTGTTACTCGCTCACATTTAA
TGTTGATGAAAGCTGGCTACAGGAAGGCCAGACGCGAATTATTTTTGATGGCGTTCC
TATTGGTTAAAAAATGAGCTGATTTAACAAAAATTTAATGCGAATTTTAACAAAATA
TTAACGTTTACAATTTAAATATTTGCTTATACAATCTTCCTGTTTTTGGGGCTTTTCTG
ATTATCAACCGGGGTACATATGATTGACATGCTAGTTTTACGATTACCGTTCATCGA
TTCTCTTGTTTGCTCCAGACTCTCA

**Supplementary Note 2 | Staple sequences triangular monomer 1**.

Triangular monomer making up the pentameric base of the DNA origami nanoshell.[3]

| Name | Sequence |
|---|---|
| side1_core_1 | AGGGGACGACGACAGTTTTTATCGGCCTCAGGAAACCGTGCATC |
| side1_core_2 | CCGAACAAGAATACCCAAAAGAACCATACATAACAGCCATGTTTTGAA |
| side1_core_3 | TTAGACGGGAGAACCCGAAGCCCTTCCTTATTTGCAGCCA |
| side1_core_4 | GAATCGATTCTACTAAGCTATATTTTCATTTAAGATTGATCAGAA |
| side1_core_5 | CCAATAATAAGAGCAAGAGCAGATAGAAACAGGGAACGTCAA |
| side1_core_6 | GTGTACAGACCAGGCGCATAGGCTATGCCACTGAGGCGCAAAACAGCT |
| side1_core_7 | ACTATTATCTGGAGCACAAACTAATAGCGCGAAACAAAGTGCTCCAT |
| side1_core_8 | AGAAAAATCTTTCATCAAGAGTAATCTTGACATTTTTGAACCGGATATTCAT |
| side1_core_9 | AAGCAAACTTAATTGCGTCTGGAAATATTTTAATTTTTTGCAATGC |
| side1_core_10 | CTTAGAGCTCCAACAGGTCAGGATTTTTTTGAGAGTAC |
| side1_core_11 | CTAGCATGCAGCAAGCCCAATAGCGAACGATCTAAAGTT |
| side1_core_12 | AGATTTGTAACACTCATAGTTAGCGTAATTATGAAACA |
| side1_core_13 | AATATTTGCATTAAATTGTTTAGACTGTTTTTATAGGCATCGTA |
| side1_core_14 | TCACGACGTTGAGCGCTAATATCCGCAAGTCAGATTGAATA |
| side1_core_15 | AGAGCCACCACCCTCAACCCTCAGATAGCTATAGCTAGCAAGGACCAT |
| side1_core_16 | AAGCCCCATTTTCAGGGATACCGTCGCCATTCAGGCTGCGCAACTG |
| side1_core_17 | TATGTTAGCCGGAGACAGTCAAATCACCATCACGCGAGTCGAAAT |
| side1_core_18 | TGATACCGCAACCTTTAATTGTATCGGTTTTTTTATCAGCTCAC |
| side1_core_19 | TTCTAAGAAGAGGACAAGAGGCAACCGCGACCTACAACGGGGCTATCA |
| side1_core_20 | TGATAAGAGGTCTTTTTTTTTGCGGATGGTCATTATA |
| side1_core_21 | CCTAATTTCAACGCTCGGGTTATATAACTATACTGTAAATAGAGAGAA |
| side1_core_22 | CATCAATGAACGGTAATCGACCATGTACCGTATCATCG |
| side1_core_23 | GCACCATTACCATTGAAAAGGTGGATTAAGCAACGGAGATCTACAAA |
| side1_core_24 | ACCAGTAAAAGTAAAACAATGCTGAACACACCCTCGGCGATC |
| side1_core_25 | GGTGCGGGCCTCTTCGTGATTGCTGGGTAATTTAACATAA |
| side1_core_26 | AATGGGAGGGATTTTGCTTTTTAACAACTTTCAAATTTTTTC |

| | |
|---|---|
| side1_core_27 | ATCATACCCTGAGAGTGATAAATGTTACTTAGGAACCGA |
| side1_core_28 | AATTTCTTAAAACGAACTAATTTTTGGAACAACATTATCCAGTCAG |
| side1_core_29 | AGGCAAGGGATAAAAATTTTTAGAACCTTTTTCATGTTTCATT |
| side1_core_30 | CAATAACGGATTCGCCCTATTACGCCAGCTGGCGAAAGG |
| side1_core_31 | CGCAGAGGGAATTAACTGAACACCAAATAGCAAACCGCCA |
| side1_core_32 | TTTCCAGATCCAGCCATCACCAGTAAACAAGAGGTCATTG |
| side1_core_33 | TTCTGTATTAGGTCACGTTGGTGTAGATGGGCCCAGGCAA |
| side1_core_34 | ACGTTGAATTTTTATCTCCAAAAAAAAGCAACCATCGCCCACGC |
| side1_core_35 | AGCTATTTTTGACAGAAATTGTGGCGTTTTATCCGGTA |
| side1_core_36 | TAATGCCGATTCAACCTGTGTAGGTAAAGATTCAAAAGGTTATTTC |
| side1_core_37 | CTTTAATTTAGTCAGAAGCTTTTTAAGCGGATTGCATACCCTG |
| side1_core_38 | GCCGGAAACGTACCCCGGTTGATATATAAGCA |
| side1_core_39 | TTTGGGAATTAGAGCCAGCAAGCCGCCACC |
| side1_core_40 | GTGAATAACCTTGCTTTGTAAATGAAATGAAACAAAATAAAGGTGGC |
| side1_core_41 | GGGAGGGAAGGTATTATCACGAAAATATGGCATG |
| side1_core_42 | GCCAGTTAATAGCAGCCTTAGACGCTGAGAAG |
| side1_core_43 | GCTTTCCGGTCAATCATATGTCCAATACTGCGGA |
| side1_core_44 | CGTCACCGACTTGGGGTCGGTTGTACCAAAAACAT |
| side1_core_45 | GATCGCACCGTTAGTACTGTAGCATTCCACAGGCGATTAT |
| side1_core_46 | CTTAGGTTAACAGTAGGGCTTAATAGCCGTTTCCAGCTACAA |
| side1_core_47 | CTGATGCATTAAAATTCTTACCAGCCGCGCCCGCCTTAAATCAAGATT |
| side1_core_48 | CAAAGAACCCTTAAGAAACGATTTATTAAGACTTTTAAGA |
| side1_core_49 | AAGGTGAAAATATTGAAAAGACAAAAGGGCGGCGGGAG |
| side1_core_50 | ACCAAGTTACCAACCTAAAACGAAGATGAACGACTGACCAACTTTGAA |
| side1_core_51 | CATGAGGAAGTTTTTTTCCATTAAATAATTTTTCGATATATTCGGT |
| side1_core_52 | GAGGTTTAGTACCGCCAAAACAGGGGGGCGCGCTTTAAAA |
| side1_core_53 | AATATGATGAGAGGGTAACGCAAGCAAAGAATTAGCAAA |
| side1_core_54 | AATAGCAAGCAAATCAGATATAGTCAAATATATCCCAATCCAAAGAT |
| side1_core_55 | CTGAGTAAGTTCTAGCGCTCCTTTATCATAAGGCCGGAAC |

| | |
|---|---|
| side1_core_56 | CCGGAATCATAATTACATTTAATGAAACTTTT |
| side1_core_57 | GCATAAAGCTAAAAGAAGCCTGTGAGAAAGGCCGATTGA |
| side1_core_58 | ACGAAGGCGCGCCGACAATGACAAGCTCCAAAAGGAGCGCAATGAATT |
| side1_core_59 | ACGGGTAAAATACGTAGGCTGACCACGTTAATGACGGTCA |
| side1_core_60 | TTATTTTCATCGTAGGAATCATTATATAAAGC |
| side1_core_61 | TATGACCCTGTAATACTTTATAAAGCCTCAGA |
| side1_core_62 | AAGGCTTAGCGAACCTCCCGACTTACATTCAACAAACGTA |
| side1_core_63 | GGAGGTCAATAACCTTTTTTTTATAGTAGTAGTTTTATTAATAGAT |
| side1_core_64 | ATTTCATTTGAATTACCTTTTTTAAGAAGATGATTCATTT |
| side2_core_1 | CCGCCACCCCTCATTAAAGCCAGAATGTTTTTAAAGATTCATTA |
| side2_core_2 | ATCGGCTGTCTTTCCTTATCATTCTTAGGCAGTAAGTCCTGTGAATTT |
| side2_core_3 | ATGGTTTAAACATATAAAAGAAACGTTTTTAAAGACAC |
| side2_core_4 | CCAGCGCCCGGAAATTCGCAGTCTCTTTTTGAATTTACCGTTCCAG |
| side2_core_5 | AGGCGGTTAGAAACCAATCAATACTAATTTACGAGCATG |
| side2_core_6 | CTCAGAATTAAGAGGGCCCGTATGTTTATCAATCCCATC |
| side2_core_7 | TTTTCCCTTTTTTAGAATCCTTGAAAACCATAGGTCTGAGAGAC |
| side2_core_8 | GTGCCAGCTGCATTAATGAATCGGCCAACGCCAGGGTGGTTTTTCTT |
| side2_core_9 | TTTTATCCTGGGTATTAAACCAAGTACCGCACTTTTTCATCGAGAACAAGCA |
| side2_core_10 | GGGGTCAGAATGCCCCAAATAAATCTCAGAGCCACCACC |
| side2_core_11 | GACTGTAGCGCGTACCGGAACCTCAGAGCGGGGAACCTATTATT |
| side2_core_12 | TGAGGCCAGTTGCTTTGTAATAACATCACGCCCCGCCAGCATTGACAGGAG |
| side2_core_13 | GTTAGAACTCAAACTACCTGAAAGC |
| side2_core_14 | TCACTGCCCGCTTTCCAGTCGGGAGTTAACGG |
| side2_core_15 | CTGTTTGACATCAGATGTCATAAACATCCCTTAGCACCGTTTAAAGAA |
| side2_core_16 | ACCACCAGAGCCGTTGATATTTGATACAGGAGTGAGTAAA |
| side2_core_17 | TAACTGATTGTTTGGATTTCAGGGCGATGGCAGCTTGACAGCGGAAT |
| side2_core_18 | TGGTGGTTACAAGAGTCCACAATCCGCCGGGC |
| side2_core_19 | AAAAGAATTAAGTACTATGCCGTACTGGTAATAAGTTTTAACTTGCGTA |
| side2_core_20 | TCACGCAATGTTTTTATAATCAGTTCACCACCCTTATAAATC |

| | |
|---|---|
| side2_core_21 | AGAGTCTGTCCATTGATTAGACGAGCGCCGCGC |
| side2_core_22 | TGAGAATCGCTTTTTATATTTAACTACCTTTTTTTTAACCTCCGG |
| side2_core_23 | AACGCCAACATGTAATCAAGAACGAATCTTACGAACAAGA |
| side2_core_24 | TTCAGCTAATGCAGAAAGTAATTCTGAAACAGAAGGATTACCACCGG |
| side2_core_25 | CACGGAATAACCGAGGAAATTTTTGCAATAATAACGAGTTACC |
| side2_core_26 | AAACAGTTTGCCTTGATAAGCGTCATACATGGCTTTTGACACAAAC |
| side2_core_27 | GACGACGACAAAGCCCGAGATAGGTTAATGCGGAGAAAGG |
| side2_core_28 | CTTTGCTCGCCGGGTTACCTGCAGCGTTGCGCCTGAGAGAGT |
| side2_core_29 | CTGCCTATTTCGCGCACAACATGTTGGGCGCGCGGGGAG |
| side2_core_30 | GTTGAGTAAAGGGCGAAAAACCGTCTAATAAACGTGGC |
| side2_core_31 | ATCAATATAATCCCTATGTTTACCAGTCCCGGAATTTGTGAGAGAT |
| side2_core_32 | AAGGAGCGCGTAACCACCACACCCACGTATAAGGCAAAATGTGAGACG |
| side2_core_33 | AGTCAATACAACGCTAACGATTTTTCGTCTTTCCAGAGTTTTGCAC |
| side2_core_34 | GATGAATATACAGTAAGCTGCAAGGCGATTAAGTTGGGTACGAAACGT |
| side2_core_35 | CTGGTCTGGTCAGCAGTAGCTCTCACGGAAAAAGAGACG |
| side2_core_36 | CCGCTACATTGTTGCCTGAGTAGAGAGGCAGGGCATTTTCGGT |
| side2_core_37 | GGGATGTCAGTACCTTTTTTTTTCATCGGGAGAAATTAATTAA |
| side2_core_38 | CTGTCCATAATGGAAGGGTTAGGGAACGGAACCAGGCGGATAAAATTGAGTT AAGC |
| side2_core_39 | CCTTGCTAAGGGAAGATTTAGCCACTACTGATTATAGACTTT |
| side2_core_40 | GGTCATAATCAAAATCTTTCATCGTCAGACGACGCCACCAGAACC |
| side2_core_41 | CAGAGCCAGGATTAGCGGGGCGCTTCTGAA |
| side2_core_42 | TCTTTTAATAGCCCCCTTATTAGCGGCCTTTAGCGTCA |
| side2_core_43 | ACAGCGCCCACCGGAAACAATCGGAAGGTGCCGTCGAGAGTATCACCG |
| side2_core_44 | CTCCGTGGTGAAGGGACAACCGCATCACCCAACGTGGACT |
| side2_core_45 | TTTGTCACAATCTTTTTATAGAAAATTCATAGTTGCTA |
| side2_core_46 | CAGATGATGGCAATTCCCAGAAGGGGGGAAAGTTTGCCA |
| side2_core_47 | CGGAACAAAGAAACCAAGCCCGGATCAAGTTTCCGGTTTTGCTCAGT |
| side2_core_48 | TCAGTAGCGACAGAAATAGGTGGGTTGATATAAGTAT |

| | |
|---|---|
| side2_core_49 | CGATAGCAGCACTTTTTGTAAAACCGCCTCTTTTTCTCAGAAATC |
| side2_core_50 | AATATAAAATTATTTGCACGTGCGATATTTTTCTTAGATTATACCGTCGCTA |
| side2_core_51 | AGAAGGACTGAGACTATATCAAAGTACCGACAAAAGGTAACGCGCCT |
| side2_core_52 | GTAAAACGACGGCCATTTTTTGCCAAGCTTTCAGGTTTTCCCAG |
| side2_core_53 | TACTCAGCCCTCAGAACAGAGAGATAATTTTTCCACACGCCAGG |
| side3_core_1 | TTGCGGGAAACGAGGGTAGCAACGAGTGAATAGATTTTAAGAACTGGC |
| side3_core_2 | TCAAATATCCAGAACGAGTAGATTAATACCGATCGTCTGAAAT |
| side3_core_3 | CAGGTCTTTCAAAAAGATTAAGAGCAGACCGGGAGATTTACCTTATGC |
| side3_core_4 | ATCGTCATAAATATTCATTCAAAAATTACCAGACAGGAATTA |
| side3_core_5 | TACAGGTAGGCTTTGACTTGCAGGGAGTTAAAGGAATTGCTATCATAA |
| side3_core_6 | TTATTAATCAATAGGAGGTAAAGTTAACGAGAGGCTTTTG |
| side3_core_7 | GCAACGACCAGTAATAACGCTCAAACGAACCAATTAGTCTTTAAT |
| side3_core_8 | CATTTTGTATCATCATATGGGTCGAGGTTTTTTCCGTTCATTTGC |
| side3_core_9 | GGTCACGCTGCGGGCGCTATTTTTGGCGCTATAGATAA |
| side3_core_10 | CCAACAGACGCCAGCCATTGCAATTTGAGTCCGAACG |
| side3_core_11 | GTAAGAAAGCCGTCAGTTGAAAGCCCGGGTAGTTTCCTGAACATACG |
| side3_core_12 | TTAATTCGATGATATCAAACCCTCAATCAATTGAGATGGAGCCTC |
| side3_core_13 | TCTAAAATATCTGTTGCGTCCGTGTTTAATTGTAGTAAATTGGGCTGCTCACAA |
| side3_core_14 | GAATCAGAGTTTTTGGGAGCTAAACAGGAGGGCAAGTGTAG |
| side3_core_15 | CCCTCAAATAAAATTCAAATTGTAAACGTTAATTTAAAAG |
| side3_core_16 | GTCAGTATTAACACCGCCTTTTTTCAACTTGTAGCAATACTTCT |
| side3_core_17 | TTCCCAATTCTGCGCAGCCCTAAAACATCGCCATT |
| side3_core_18 | ACATTTCCCATAAATGAATCCCAAAAGAGTTAAAT |
| side3_core_19 | CGCCAAAGACGATAAAAACCAAAATAGACGCAGAAAAC |
| side3_core_20 | CAGAACGATCAACTTTAATCATTGAGCTCAACAGCTTCAA |
| side3_core_21 | GTCGGGGTCATTGCAGGCGTTTTTTTTCGCACTCTACGGTGGTG |
| side3_core_22 | ACAGGCGGCGCGGTCCTACATTTGTAGATTAGTACGTGGC |
| side3_core_23 | TCAACGTAACAAAGCTGCTCATTCGCTACAGA |
| side3_core_24 | TGTACATCGACATAATTTTTAAAATCCCGTAAAACGCCAGCAGT |

| | |
|---|---|
| side3_core_25 | TGCATCAGACGATCCATGTAAAGCGGTCCACGAATCATGG |
| side3_core_26 | GCGGTTGCCTGGTTTGCCCCTTTTGCAGGCGAAAATCGCCTGGCC |
| side3_core_27 | TGTTCTGCAGATACATAAGAAAGACTGAGAATGA |
| side3_core_28 | TGCCGTTCCGGCAAACCTTTAGTTCGACAACTCGTAGCACTAAATCGGA |
| side3_core_29 | AACAACCCGTCGGATTCTCCGTGAGAATAGACAGAGGGGCCCTCGTT |
| side3_core_30 | CAGCTTTCATCAACATTAAATGTGAGCGAGTCAGCTCAT |
| side3_core_31 | CAGAAACGACTTGTAGAATTTTTGTCAGCGTGGTGCCATCCCA |
| side3_core_32 | GAATAATACAGTTTCAGCGGAGTGGGAACAAACGGCGGATTGACCGT |
| side3_core_33 | CCAGCGGTGCTTTTTGGTGCCCCCGGTATTTTTTGGGTAAAGGTTT |
| side3_core_34 | CGCAACCATTTTTCTTACGGCTGGAGGTCTGTTGCCCTGCGGCT |
| side3_core_35 | CGCTGAGGGGACTAAAGACTTTTTTACCCAAAGACGTTGGGA |
| side3_core_36 | TGAGGATCGAATTGAGAGTTGGCAATGAAAAATCTAAAGCATCACCTAACAG AG |
| side3_core_37 | CTCACAGTGTTTCTGCACAACTAAAGGATTTA |
| side3_core_38 | AGTCACAATTTATTTACATTGGCAGACGCTCATGGAAA |
| side3_core_39 | TATTTTTATTCTGGGAAGTATTAGACGTTATGCTGATCGTGCC |
| side3_core_40 | TGCGGCGGCCGGGTCAGTCCAGCATCAGCTCGATAACGGAACGTGCCG |
| side3_core_41 | TTTTTAACTATTTTGTTGCTTTAAATTCACC |
| side3_core_42 | TGCAGCAAGCCTGGGGTGCCTAATGAGTGAGCTTTTTAACTCACATTAATTG |
| side3_core_43 | TGTGAAATTGTTATCCATCTGGTCGAAGGTTAGTGAGGCGACAGACAA |
| side3_core_44 | AGCCGGAAGCATAAAGGCGCAGTGGAATTCGTGGTATGAGGCCGTTTT |
| side3_core_45 | CCTGTAGCCAAAAATAATTCGCGTAAATTAAATCCTTTGCAACATTAT |
| side3_core_46 | GAAAGATTAGTAAGAGCAATGCTTTCGAGGTG |
| side3_core_47 | CAGGAAAAACATTTACAAACAATGATGAAGACGCCAT |
| side3_core_48 | CTGAGAAGATTAACCGAGTGCCACGTTTTTTGAGAGCC |
| side3_core_49 | AAGGGATTTTAGTTTTTCAGGAACGGTACGCCTTCACC |
| side3_core_50 | CATATCCAGAACAATTTTTTTTACGATAGAACCTTTTTTTCTGATCGG |
| side3_core_51 | TACCTACATTTTGAAAGGGACGAATGGCTCGCTTAGGAGCACTACAGCACG |
| side3_core_52 | AAATAGTTTGACCATCATCCAATAA |

| | |
|---|---|
| side3_core_53 | TATGCAACTAACAGTTGAAGCGAACGAAGCCC |
| side3_core_54 | AGCAGCAAAATCAACAGCCGATTATCATAGCTCCGAGCTCTCACTGCG |
| side3_core_55 | GCGCGAACTGATAGAAGATAATGCTGAACCTCAATTTTAAA |
| side3_core_56 | CCATATAAAGTACGGTTGAATATAATGCTGT |
| side1_hub_1 | CGTTATACATTTCATCTTCTGACCTAATAGAAAAATCGCAAGACAATTACC |
| side1_hub_2 | GCGAACACTCATCTTTAATAAACAAACATCAAGAAAACAAAATTA |
| side1_hub_3 | AAGAATACACTAAAGAGTTTGAAATACCGACCGTGTGATAAATAA |
| side1_hub_4 | AATCAATATGAGCAAAATGGAAACCAAAATCGTTGGGAAG |
| side1_hub_5 | ATTATTTATTTTAGAATCCAAGCCTGTTTAGTATCATATG |
| side1_hub_6 | GGCGTTAAATAAGGACCCCCAACAGCCCTGAGTTTCG |
| side1_hub_7 | TTTGTTTAAGCGCATATGTGACCAAGTTAAGTACATA |
| side1_hub_8 | ATTACATTTAACATTGTCGTCAGCGCCATCTTCTGGT |
| side2_hole_1 | TGGGCGGTTGATCAAGTTTTTTGTCCGTGAACCAAGA |
| side2_hole_2 | CCAGTAATAAGAGATCAAAATATAAAAACAGAAATAA |
| side2_hole_3 | GTAACAGTAAGTTTATAAAATAATACAATAGAAGGCATTTTCGAG |
| side2_hole_4 | ACCCTAAAGGGAGCCCCCGAAAGCGACCAACGTCGTTGTTCCGTG |
| side2_hole_5 | TTCAGGTTCCGCCACGAACCTACCCCTCAAGATGAAAGTA |
| side2_hole_6 | AGGTGGAGTAACGTCAAGAAATTGCGTAGATT |
| side2_hole_7 | ATGCCAACGGCACACTGGTAGTTTGGACCGAAATCCGTGCTTT |
| side2_hole_8 | TTCAGCAAATCAACCTGTCGGCAACAGCTGATTGCCCAGAATCCCTCGTTA |
| side3_hole_1 | TGAATTAGGAATACAGCATCGGTCGTCACCCTCAGCAG |
| side3_hole_2 | TGCCAGTTTGGTAATAGTAAAATTTTAGTTTTGCAAG |
| side3_hole_3 | AAAGCCGCACATCCTCTCGCTGGCAGCCTCCGGGTGCTGCTACCGGGG |
| side3_hole_4 | AGCGGATCAAACTTAAATTTCTGCCTGGCCTTGCCAGAGC |
| side3_hole_5 | AGGCTTGCCCTGACGAGAAACACCGAAAGACCACATTCAACTAATTC |
| side3_hole_6 | GAACAACTAAAGGCCGCTTCGAGGCATCATCAGTT |
| side3_hole_7 | GTTTTTTCCACGGTCAGGCCAGAACGCCTGTGCACTCTGTTTCCACAC |

**Supplementary Note 3 | Staple sequences triangular monomer 2**.

Triangular monomer making up the protruding spikes of the DNA origami nanoshell.[3]

| **Name** | **Sequence** |
|---|---|
| side1_core_1 | GGAGGTGTGGTTGCGGACGCAGAAAACGGATAGTTGGGTA |
| side1_core_2 | GTCAAAGGCAGTTTGGGTAGAACGGTAGGGGGTTTCTGCC |
| side1_core_3 | CCACAAGATTAAGCAAATCAGATAGAGGCGTTTTTTTAGCGA |
| side1_core_4 | CTCAGAGCATAAAGCTTAAGAAAAGTAAGCAGATAGCCG |
| side1_core_5 | CTCCGGCTTAGGTTTTTTGGGTTATATAACGAATTATC |
| side1_core_6 | ATCCCATCATCGGCTGACCGACAATTAGGCAGAGGCATT |
| side1_core_7 | GAGCACATCCTCATAACGACCGCAAGAGCCGCACCAGTTGGG |
| side1_core_8 | AGAGAGAAATTGAGTTAAGCCCAAGAGATAAC |
| side1_core_9 | AAATCAAGATAATTATTCATTTTTTTCAATAACATAAGTCAGAGG |
| side1_core_10 | TCATTGCAAGTCTCTGTGGTGCTGCGGCCAGAATGCCAT |
| side1_core_11 | ATTAACACCGCTTCTGCTCATTTGCAGCGGGGGTCTGGTC |
| side1_core_12 | CCAAAAGAACAACGCGGTCCGTTAAGGATTGCCGTGTACCA |
| side1_core_13 | GGCATTCCAAGAACGGGTCAGTACCATCACCCAAATC |
| side1_core_14 | ATGCGTTATACAAATAGAACGCTAGAAGGCTTATCCG |
| side1_core_15 | AATCAATACTAATTTAAATGCAGAACGCGCCTGTTTATCATAAAGT |
| side1_core_16 | AAAACGACGTTGGCAAATCAACAGAATCAATATCTGGTCA |
| side1_core_17 | GTAATTGAAAGTTTTTCAAGCAAGACCAAGTA |
| side1_core_18 | TTAACGTCTTTTTAAAATGAAGGGAGCCCCCTTTTTATTTAGAGCTTGGTTTTTAT |
| side1_core_19 | CTCAACAATTTTCATCGTAGCGCTACGTGA |
| side1_core_20 | TCACGGTCATACCTAAAGCCAACGTTCGAGCC |
| side1_core_21 | GCCGTTTAGCAGCAGAACGTGCGATGCTAACGTGG |
| side1_core_22 | AGCTATCTTACCGAAGCCCTTTTAAATCGGT |
| side1_core_23 | GGGTTGAGTGTTGTTCGTATTCTATCTTACCATCAGGAATCATTACC |
| side1_core_24 | TATATGTAGAATTTATCAAAATCATTTTTTGGTCTGAG |
| side1_core_25 | GCGCTAATGCGCCCAATAGCACCTGAGCAAAAGA |

| | |
|---|---|
| side1_core_26 | AGATGATGGCTCTTTAGGAGCACTAACAACTATTTTTAGATTAGAGCCGTC |
| side1_core_27 | ATCCTATCAGGGCGATGGCGGGTAAAGTTAAACCGGACTTAACAAGAG |
| side1_core_28 | ACGCCAACCGCACTCTAGAAACCTGAAAAATAAACCCTC |
| side1_core_29 | CATGTAATAAGGTAAAGTAATTCTGTCTTTTTAGACTTTCATCT |
| side1_core_30 | CCTGAGAGATCAAAAATAATTCGCCGCCAGCTCGCCATGTTT |
| side1_core_31 | TTCGCACTGTTTCCTGAACAAGAAAAATAATGGCCAGT |
| side1_core_32 | TTTTGAATGGTTTTTTATTAGTCTAGTATTTTTAAGAACTCAAACT |
| side1_core_33 | TTCCGGCAGTAAAAAAAATGCCAATTACGGCTAGCTGTT |
| side1_core_34 | GGGATAGCGGAACGCCTCTGGAGCAAACAAGAAAAGGCCG |
| side1_core_35 | TCTCACGGTACATCGACATGGAGAGGGTAGCT |
| side1_core_36 | GCGCGCCTGTGCAAATAAGAGAATAACAATAGATAAGACCTGCAGCCA |
| side1_core_37 | AGACTACCGAGTGAATAACTTTTTTTGCTTCTGTAAAAATCAA |
| side1_core_38 | TCTTTCCTTATCGCTCAACAGTGGCCAAGCTACGTTGT |
| side1_core_39 | CCACGCTGAGAGCCAGGTGAGGCGGTATTAACCGTTTTTGTAGGGCT |
| side1_core_40 | TCACCTTGCCGAACGAACCACCAGAGGACGCAAATTAACCGTTG |
| side1_core_41 | AACAACATGTTCAGCTCGAGCATGTTTTTAACCAAATATCCTAAAGCA |
| side1_core_42 | GCGGTGCTGTCACTCGGGCGCCCAGCATCCGCCAG |
| side1_core_43 | CGGTGCCCTCGTTAACGGCATCACCACGGGACAGCGGTTTGTTA |
| side1_core_44 | TTAATGCGCGAACTGACTAAAATAAATTCATCCTGAACCT |
| side1_core_45 | AATATACACGGGAGAATTAACTGTTTTTACACCCTGAACAAAAAACAGGGAA |
| side1_core_46 | AATAAATAAATATAATCCTGTTTTTTTGTTTGGATTATATCATATT |
| side1_core_47 | AGCCTGTTTAGTTTTTTTCATTAATTGAGATTTTTTCGCCAAATA |
| side1_core_48 | TATATGTATCGAGAATGGGGTCGCAGAAGATAAAACAGAGCAGCAAA |
| side1_core_49 | AATCAGTGTTTTTGGCCACCGAGTAAAAAAACATCACTTGCCTG |
| side1_core_50 | GGCGAAAGGGGGATGTGCTGCAAGAACCAATA |
| side1_core_51 | ACGCCAGGGTTTTCCTGATAATCATCAAACTTAAATCTG |
| side2_core_1 | TTTTTCATAGGTTTAGGCCCGTATAAACAGTTTTGACAGGTCTTTGAC |
| side2_core_2 | TTTGAAAGAGTTAACCCTCGTTTACCAGACGACGAT |
| side2_core_3 | ACCGTCGCCCTGAGATAGCATTACGGCGGATTGACCGTAAGTTTGAG |

| | |
|---|---|
| side2_core_4 | CACTACGGGGTTGCCCTGATAGCTGCATTAATGAATCGGCCTAACCGA |
| side2_core_5 | TTCCATTATAAATTGGGTCAGGACACAGGTAGAGGTCTTTA |
| side2_core_6 | CCACGCAGTGCCGGAAACCAGGCTCCGGCACCGCTTCTG |
| side2_core_7 | GATACCGATAGTTGCGCCGACAAAGGCTGAGTAATGC |
| side2_core_8 | CTTGCTTTCGAGGTGAATTTCTTATACTCAGG |
| side2_core_9 | GAACAAAACATCCAATAAATCTGATATTTTTATTAATGCCAAAGAGACAGT |
| side2_core_10 | AGATAAGGCACCAACCTCTGCTCATGTGTACAGAGCAACACTATCATGAGG |
| side2_core_11 | TACAAGCTGATGAACGTCAGTGAAACACAACTAATCGCCAAAAGGAA |
| side2_core_12 | ATTCTACTAAAATACACGAAAATCCTGTTCAGCCTCCGGCCA |
| side2_core_13 | CGGAACAAGATTTACACCAGAAACAAAGAAAACGAAAGCGCG |
| side2_core_14 | ATCAGTTGACATTATTGTTGGGAAGAAAAATGGGAGTT |
| side2_core_15 | GTCTGGCCTTTTTTTCTGTAGCCAAAGGTTTTTTATCAGGTCATTG |
| side2_core_16 | GCTTTCATCAACATTAGCGCAACTGAATTTGTGGAAGATC |
| side2_core_17 | GGAATACCACATTTACGAGCCGGAACCTGTCGTGCCAAAACGAACTAA |
| side2_core_18 | AAAATCCCTGGCATGATTAAGACTTTTTCCTTATTACGCAGTAACGGAATAC |
| side2_core_19 | TACTCATTTGGGGCGCGACGGAGATTTGTATTGACCAACTTAGTTTG |
| side2_core_20 | ATGTTAGCAAACGTAGAATAGTAGGAGTTGCAGCCCTTCA |
| side2_core_21 | TTACGGGAATCAACGTACGAGTAGAACGGGTAAAAGGCCG |
| side2_core_22 | TGAAATTGGCCTGGGGTGCCTAATGAGTTTTTGAGCAGACGATCCAGCGCAG |
| side2_core_23 | GCAAAATCTTAGCTATATTTATAACCACGG |
| side2_core_24 | TAAAGGTGAAAATCCGCGACCTGCATTGATAAATCCGCCTCC |
| side2_core_25 | TCCTGTGCCGCCTGGGTATTGGGGGGACGACGCCAGCTT |
| side2_core_26 | ATCCCCTCCACACATAAGAGACGGGCAACGGTCACGGGCATCA |
| side2_core_27 | GGGTACCTCCACGCTGGTTACAGAAAAGGTTTGGTGT |
| side2_core_28 | GTAACAGTTACCGCCACCCTCAGATTTATCAGGACAGCATCG |
| side2_core_29 | AACCGTGCATCTGCCATGGGATAAGCTGATTGCAAGCGG |
| side2_core_30 | CGGTTTGCAGGTTTCTGCACTCCAGACAGTATCGGCCTCATCTCCGTGG |
| side2_core_31 | ATTTTGATGAGAGATAGACTTTTTTTCTCCGTGGTGAAACGTACAG |
| side2_core_32 | GAGCTCGATTCGCGTCCGTGAGCATCAAAAGAAATCG |

| | |
|---|---|
| side2_core_33 | TCAGCAAACCTGCATCTAACTCACATTTTTTAATTGCGTTGCGCTC |
| side2_core_34 | CATCAATATTTTTGATATTCAACCGTTCTCGTGAGAGATCTACA |
| side2_core_35 | AGCACGCGTGCCTTTTTGTTCATTCGTAATTTTTTATGGTGGCGG |
| side2_core_36 | TCCCAATTCTGCGAACCCTTATAACTCCTCACAGGTGCCCCAGCAGG |
| side2_core_37 | CGAGATATCCACTATTATTTTCGTCTCTTTTTTCGCGCAATAAT |
| side2_core_38 | GAGGAAGTCACTAAAACACAAGCGTCATACAT |
| side2_core_39 | TGGTGTGTGGGTCACTGTTGCCCTGTTTTTGGCTGGTA |
| side2_core_40 | TTGCTCGTCATATTTTTACATCCCTTACACTCGGCGAA |
| side2_core_41 | ACGGTCAATCATAAGGGAGCATAGGCATTATACCAAGAGGCA |
| side2_core_42 | ACCGAACCATCGCCTTCGCAAATAGAAAATTCATATGGTTTACC |
| side2_core_43 | AACGCGCGTGGTTTTTAGTGTAAATTATCCGCTCACAAT |
| side2_core_44 | ACCAGTCCCGGTTGGGAAGGGCGATCGGTGCGTTTTTGCCTCTTCGCTATTA |
| side2_core_45 | CTTTTCACCAGTGGCCGCATCGTTATATTCGACCATCGC |
| side2_core_46 | GTCATAGCTGTGTCGACCCCAGCGTGGCTGACTTACCCAATAGCGTC |
| side2_core_47 | TTCATCGGCATTTTCGAGCGCCAAAGACAAAAGGGCGAC |
| side2_core_48 | ATGGGTAAGTGGTGCCATCTTTTTCACGCAACCAGCCGGCAGC |
| side2_core_49 | CGCCAGGGGGGAGAGGACTGCCCGCTTTCCAGTCGGGAAAGCATAA |
| side2_core_50 | AACAAAGATTAGCAAAATTTTTTAAGCAATAAAGCCAAATCAC |
| side2_core_51 | GCAAAGACAAGGTGGCAACATATATGGTGATGGTGGTTCCGAATAGCC |
| side3_core_1 | TAATTTGCCAGTTACAAAATAAAAAGGAGCGTTAGATTCGCCTGATT |
| side3_core_2 | GGTAAATAGCGTCTTTCCGTAATCGCCAGCAAACCATCGATTTGCGGATGCTC CTT |
| side3_core_3 | ATTCAACTTTAGCGTCAGTTTTTCTGTAGCGCGTTGCAGGTCA |
| side3_core_4 | GAGAAAGGACAGGAACGGTACGCAACAATATTTTTCAGG |
| side3_core_5 | ACAAATAACTCTGAATTTATTTTTCGTTCCAGTTCAAGGTTGAG |
| side3_core_6 | GCTAACGACCGTTTTAAATATGCACATATAAC |
| side3_core_7 | TTTGCACCCAGCTACAGAGGTTTTAATTACATAAAAATTACTAGAAAA |
| side3_core_8 | GCGCATTAGATCGCGCAGAGGCGTAGATACCAAG |
| side3_core_9 | GACGATTGTTTTTCCTTGATATTCACAACTGGTAATAAGTTTTA |

| | |
|---|---|
| side3_core_10 | ATCGGCCTGTGGCACAGACAATATAATAGATACCTGATTATC |
| side3_core_11 | TACCGCCAAGAACCCTTCTGACCTTAGACTTTACCACCAGAAGGAGCG |
| side3_core_12 | AAAACGCTCTATAAAACAGAAATACCTTAGAATGAATTAC |
| side3_core_13 | CAAATAAGAACGTGGCTTACAAAAGTAACAGTCAGTACATATCGTCGCTTTAACA |
| side3_core_14 | CACCCTCAGAGCCACCAAATCTCCAGCAACGGACAACTTT |
| side3_core_15 | TTATCACCGTCACCGTTTTTCTTGAGCCATTTGGAATTATTCAT |
| side3_core_16 | GAATTAGAAGTAGCGATAAAGCCATAGCAGCAAGTTTCGTCACAGACAGTAAATGA |
| side3_core_17 | AATCACCAGTAGCTCAACATGGAACGAGGCGCAG |
| side3_core_18 | ATAGCAAGCCCAATAGAGGAATTGGTGAGAATAGAAAGGA |
| side3_core_19 | ACCGCCACCCTTTTTCAGAACCGCACGGTTTTTGTCAGTGCCTTGA |
| side3_core_20 | ATACATTTGAGGATTTAGAAGTATGAAAGCGT |
| side3_core_21 | ACAAACAATTCGACAACTCGTATCTGGCCAAAATTATTTGCACGAAG |
| side3_core_22 | AGCGCAGTATCCTCATCAGAATCAAGTTTGCCCGATTGAGGGAGGGAA |
| side3_core_23 | ATAACGGCAATTTCATTTCCTTGAACCTCCGACCGTGTGAT |
| side3_core_24 | ACGTTATACAACTAAGAACCCATAGACGTTAGCCCTCATCCTTTAAT |
| side3_core_25 | AGCAATACATGTGAGAAGTACGGGGAAAGCCGGCGAAACGATTTTTTGT |
| side3_core_26 | AATTTTTTCACGTTGAACCCTCATTTTGCTAAATGATACAAACGCCTG |
| side3_core_27 | TATGGGATTTTCAGGGCAAACTACGGAGTGTA |
| side3_core_28 | ATAATGGATTTAACGTCAGTTCTTTGATTAGT |
| side3_core_29 | TTTTTCAAAATTTAAGACGCTAATTTTCAAGAAATCATCGGG |
| side3_core_30 | CCTGACTATTATAGTCAGCTTCAATGAATTACCAACAGTT |
| side3_core_31 | GGCTTTTGCTACAGAGGCTTTTTTTGAGGACTAAAGACCAGCGAAA |
| side3_core_32 | GTAACGATCCAGTCACACGACCGTAACACTGAGCGGTCCCAATGAA |
| side3_core_33 | TCTGACCTAATATATTTGCAAATCCAATCGC |
| side3_core_34 | GAAATACAAATGCTTTAAAAGATTAAGAGGCGCGAGAAAAC |
| side3_core_35 | AAATAAGGCGTTATATATTAATTGAGAAGAGAACCTACCACAAAGAA |
| side3_core_36 | GCGGAATCGAGAATGACCATAAATCAATTTTTAATCAAAGATTC |
| side3_core_37 | CAATACTTTGATAAGAGCGAACCATTTTCTGTCAGCGGACGAATAAT |

| | |
|---|---|
| side3_core_38 | AGAATAACTTTTTTCAAGAAAGCTTTGATTGCTATTTATTTATCCCAATC |
| side3_core_39 | ACACCGGAATCATAGAAGTTTTGCTGAATCCC |
| side3_core_40 | CAGAGGGAGCTTAATTGCTACAAGGCCGGA |
| side3_core_41 | GCTTTTGCTTAGAATATAATGCTGTAGCACCTGAATC |
| side3_core_42 | ACAAAATTGAAGCCTTACCTCCCGACTTGCGGCTGGAAGT |
| side3_core_43 | AAAATAAAATGTTTTTTTAGACTAGGCATAGTAAGACCAGGC |
| side3_core_44 | ATGCGCCGCTACAGGGAACGTGCTGGAGGCCG |
| side3_core_45 | AGGGTTAGTCAATAGTAATGCTGATTAGTTAAGACGACAAT |
| side3_core_46 | AATCATTGATCTTGACAAGTTTTTACCGGATATTCACTTCATC |
| side3_core_47 | ACTTCTGAAAGAATACTGCTGGTAATATCCAGCAGAATCC |
| side3_core_48 | AGTTGATACCATTAGATACTCCATGTTATTTTTTTAGTTGACGGA |
| side3_core_49 | TACGGTGTATTTTATCCATTACCAGGCGCTAGGGCGCTGCGCGCTTA |
| side3_core_50 | GAACGAGGGTAAAAAAAAGGCTCCAAAAGGAGTTTTTCTTTAATTGTATCGG |
| side3_core_51 | GAGTAACATTATCATTAAGACAAATTAGATTAATGGTTT |
| side3_core_52 | AGATGATGAAACAAACATAATGGAAAACCTTTTATGCGTAGA |
| side3_core_53 | GCGATAGCGAAAAGCCCGAAAGACTTCAAATTAATTTT |
| side3_core_54 | TGGCTTAGGGTAATAGCCAAAATAGCGAGAGTTCATTCACTAAAG |
| side3_core_55 | AAGAGTAAGGTCATTGAATGGAATAGCATTCCACCAGTA |
| side3_core_56 | TTATACCAGCTTGAGATGGTTTAATTTTTTTCAACTTT |
| side3_core_57 | AACGATTAGAGAGTAAGTTAGCAACGTCAGAGCGGGAG |
| side3_core_58 | CTTATGCGATTTTTTTTAAGAACTGGCTCAACCCTCAG |
| side3_core_59 | CCAACAGGTCAGATATTTTGTCGAAAAGTTTTTGCCCGA |
| side3_core_60 | ATCGCGTTAGCAAACTCAGAAAACGTCATAAATATTCAT |
| side3_core_61 | AGACCGGATTAATTCGAGAAGCAAAGCGGATTGCATCAAAACAGTT |
| side1_hole_1 | TTGAAAGGAATTGAGGAAGGTTATTAGCCCTAAAA |
| side1_hole_2 | GCGAAAAACCGTAGATAATAAGAGCAAGAAACAATGAATATTTTAAATG |
| side1_hole_3 | AAACAGGAAGATTAACTAGCATGTCAATCATA |
| side1_hole_4 | CATCGCCATTAGAGTCTGTCCATCTGCCGTAA |
| side1_hole_5 | TGTACCCCGGTCAGTCACGTTCAGAGGTGGAGCCGGATGCCGGCAATCCGC |

| | |
|---|---|
| side1_hole_6 | AGCACTAAATCGGAACCCTAAAAATAGCAGCCTTTAC |
| side1_hole_7 | CAATGCCTGAGTAATGTGTCTTTAGTGATGAACCATGGTGCTG |
| side1_hub_1 | GATTCAAATACTTTTGAAATATTTAAATTGTAAACGT |
| side1_hub_2 | TAATATTTTGTTACATTTTTTGCGATTAAACCTCACCGGAAACAA |
| side1_hub_3 | TAAAGGGTGAGGAATCGATCGGTTGTGAAAAAGAGTATGAGCC |
| side1_hub_4 | GAACGGTAAATCAGCTAAATTCGCATTAAATTAGAAAAGCCCCAA |
| side1_hub_5 | ATCGTAAGTATAAGCCGGGAGAAGCCTTTATTTCAA |
| side1_hub_6 | CGCAAGGATAAAACCCTCATAATAGCAATACTCCAAC |
| side1_hub_7 | AGGCGGCAGGTAAAAAATAGAAATTTTACATTATGACCCTG |
| side2_hole_1 | GCCACCACCCTCAGAGCCGCCTCTGAAACATG |
| side2_hole_2 | CAAGGCAAAGATTACCAGAAGGAAACCGAGGAAACAAAGAAAC |
| side2_hole_3 | AATAAGTTTATTTTGTCTCAGAACCGCCACCCTCAGA |
| side2_hole_4 | GTAACAACCCGTAGCATACAGG |
| side2_hole_5 | AAAGTATTAAGTGACAACAGTCGCTGAGGCTTGCACTACGTTAACGAGAAA |
| side2_hole_6 | AAAGCGCCATTCGCCATTCAGGCTAATGTGAGCGA |
| side2_hub_1 | AGGATTAGGATTACCTATTATACCAGAACAAACAAAG |
| side2_hub_2 | AAATACGAGCCCGGTTCGGAAGCGGGGTTAAATCACCGGA |
| side2_hub_3 | ACCCCGCCACCCACAATCAATGGTCAATAACCTGTGAGTAGAT |
| side2_hub_4 | CTGCCTATAATAGGTGGGGTTGATATAAGTATACTCCTCAAGAGA |
| side2_hub_5 | AGTGCCGTCGAGATATCACCGAACAGCTTCTTTTGCGGGATCGTC |
| side2_hub_6 | AGAGCCGCCATAATCATTGCTCAGTACCAGGCGGATA |
| side2_hub_7 | TTTGCCATCTTTTCGCCAGCAAATGCCCCAAAGAATA |
| side2_hub_8 | CACCACCCTCAGAGAGAGCCACCACCGGAACCCCTTATTAGCG |
| side3_hub_1 | GATTTTAGAAGGGAAATGGTTGC |
| side3_hub_2 | CTAAACATTCCTCGTTAGAATCAACGCTGCGCGTAA |
| side3_hub_3 | CAGATTCACATGGAAAGGATTATTGGGACATTTAAATCCT |
| side3_hub_4 | CTCAATCGTCTGAAATTACCTACACAACAGGAATTAAAGGAGAAACA |
| side3_hub_5 | TTTGACGAGCACGTATCGCGTACTGAAAGCGACAGCCATA |
| side3_hub_6 | TTGCGGAATATCAACAGAGATGCCATTGTTTTGACG |

| | |
|---|---|
| side3_hub_7 | CCACCACACCCGCGCAAGTGTCCAGAGCCTTACCAAC |
| side3_hub_8 | TCTTTCCGTACCAGTAATAAAATACATTGG |
